# Synaptic imbalance and increased inhibition impair motor function in SMA

**DOI:** 10.1101/2024.08.30.610545

**Authors:** Emily V. Fletcher, Joshua I. Chalif, Travis M. Rotterman, John G. Pagiazitis, Meaghan Van Alstyne, Nandhini Sivakumar, Joseph E. Rabinowitz, Livio Pellizzoni, Francisco J. Alvarez, George Z. Mentis

## Abstract

Movement is executed through the balanced action of excitatory and inhibitory neurotransmission in motor circuits of the spinal cord. Short-term perturbations in one of the two types of transmission are counteracted by homeostatic changes of the opposing type. Prolonged failure to balance excitatory and inhibitory drive results in dysfunction at the single neuron, as well as neuronal network levels. However, whether dysfunction in one or both types of neurotransmission leads to pathogenicity in neurodegenerative diseases characterized by select synaptic deficits is not known. Here, we used mouse genetics, functional assays, morphological methods, and viral-mediated approaches to uncover the pathogenic contribution of unbalanced excitation-inhibition neurotransmission in a mouse model of spinal muscular atrophy (SMA). We show that vulnerable motor circuits in the SMA spinal cord fail to respond homeostatically to the reduction of excitatory drive and instead increase inhibition. This imposes an excessive burden on motor neurons and further restricts their recruitment to activate muscle contraction. Importantly, genetic or pharmacological reduction of inhibitory synaptic drive improves neuronal function and provides behavioural benefit in SMA mice. Our findings identify the lack of excitation-inhibition homeostasis as a major maladaptive mechanism in SMA, by which the combined effects of reduced excitation and increased inhibition diminish the capacity of premotor commands to recruit motor neurons and elicit muscle contractions.

## INTRODUCTION

The balance between excitatory and inhibitory synaptic transmission is essential for normal neuronal function at the cellular and the neuronal circuit level (He and Cline, 2019). Failure to maintain a balanced excitation-inhibition drive has been proposed to lead to neuronal network dysfunction, as observed in several neurological diseases, including autism (Rubenstein and Merzenich, 2003), schizophrenia (Kehrer et al., 2008), fragile X (Gibson et al., 2008), Rett (Dani et al., 2005) and Angelman syndromes (Wallace et al., 2012). Under healthy conditions, individual neurons and neuronal circuits adjust the balance between excitation and inhibition following perturbation of incoming synaptic activity (He et al., 2016; Zhou et al., 2014). However, reduced excitatory neurotransmission has been reported to decrease inhibitory currents and synaptic puncta (He et al., 2018). In contrast, reduction in inhibitory inputs did not alter excitatory inputs (Shen et al., 2011), suggesting that maintenance of excitation-inhibition balance is not an automatic response, and excitatory neurotransmission likely exerts a dominant role. Although the mechanisms responsible for the regulation of excitation-inhibition balance are slowly emerging, it is currently unclear whether changes in one or both types of neurotransmission contribute to pathology in neurodegenerative disease.

It has been proposed that mutations during early development may specifically affect the GABAergic system, resulting in excitation-inhibition imbalance leading to pathological overexcitation of neurons (Nelson and Valakh, 2015). Whether neuronal overexcitation is involved in diseases affecting motor control and movement has been highly debated (Delestrée et al., 2014; Jensen et al., 2020; 2021; Manuel, 2021; Manuel and Zytnicki, 2021; Martínez-Silva et al., 2018). In amyotrophic lateral sclerosis (ALS) whether the motor neuron disease results from hyper- or hypo-excitability is hotly contested. Nevertheless, there is evidence from clinical studies and mouse models pointing towards an essential role of pathological disinhibition in motor cortex and spinal motor neurons during disease progression, more so perhaps than glutamatergic excitation (Gelon et al., 2022; Scamps et al., 2021; Turner and Kiernan, 2012). Recent studies in ALS models highlighted a synaptic deficit from a major premotor inhibitory input that originates from V1 interneurons which, when counteracted, ameliorates disease pathology (Allodi et al., 2021; Cavarsan et al., 2023; Montañana-Rosell et al., 2024; Mora et al., 2024; Salamatina et al., 2020). Here we sought to address this issue by investigating a potential role for inhibitory dysfunction in the neurodegenerative disease spinal muscular atrophy (SMA).

SMA is caused by deletion or mutation of the *Survival Motor Neuron 1* (SMN1) gene, leading to ubiquitous severe deficiency of the SMN protein (Lefebvre et al., 1995; Lefebvre et al., 1997; Tisdale and Pellizzoni, 2015). The hallmarks of disease in patients and mouse models are select death of motor neurons, muscle atrophy and severe reduction of spinal reflexes (Tisdale and Pellizzoni, 2015). Importantly, work from our lab and that of others have demonstrated that SMA is a disease of motor circuits (Fletcher et al., 2017; Imlach et al., 2012; Ling et al., 2010; Lotti et al., 2012; Mentis et al., 2011; Shorrock et al., 2018). Vulnerable SMA motor neurons receive less excitatory drive from proprioceptive glutamatergic synapses (Fletcher *et al*., 2017; Simon et al., 2019) as well as other excitatory premotor interneurons (Ling *et al*., 2010; Simon et al., 2016). However, whether the inhibitory synaptic drive on motor neurons in SMA mice is affected is not known. To address this critical question, here we studied inhibitory synapses on SMA motor neurons originated from two major classes of V1 inhibitory interneurons that tightly modulate motor neuron firing: Renshaw cells and Ia reciprocal inhibitory interneurons (Alvarez et al., 2013; Alvarez and Fyffe, 2007; Bhumbra et al., 2014; Geertsen et al., 2011; Hultborn et al., 2004; Hultborn and Pierrot-Deseilligny, 1979; Mentis et al., 2005; Moore et al., 2015; Sweeney et al., 2018). Using mouse genetics, physiological, morphological, and behavioural assays we found that vulnerable motor neurons in SMA mouse models exhibit an unexpectedly higher density of GABAergic and glycinergic synapses which impose an excessive inhibitory burden on motor neurons and their recruitment. This unwarranted excessive inhibition on vulnerable SMA motor neurons is likely the result of a maladaptive response to the initial reduction of glutamatergic excitatory drive and contributes to motor dysfunction in SMA. Importantly, counteracting excessive inhibition provides behavioural benefit in SMA mice, suggesting a novel avenue of potential therapeutic intervention.

## RESULTS

### Reduced activation of Renshaw cells by proprioceptive synapses at the onset of SMA

Vulnerable motor neurons in the first or second lumbar segment (L1/2) innervating proximal or axial musculature, receive reduced activation from proprioceptive synapses in a severe mouse model of SMA (Fletcher *et al*., 2017; Mentis *et al*., 2011). The reduction in proprioceptive-mediated synaptic transmission is initially due to the impairment of glutamate release (Fletcher *et al*., 2017) followed by the elimination of synapses at later stages of disease (Fletcher *et al*., 2017). Diminished glutamatergic transmission from proprioceptive synapses alters the electrophysiological properties of motor neurons and repetitive firing (Fletcher *et al*., 2017). Whether inhibitory synapses also modulate firing of SMA motor neurons is however unknown. It is also unclear whether any homeostatic mechanisms operate on the inhibitory inputs of motor neurons to counterbalance excitatory synapse dysfunction. To address these questions, we investigated inhibitory synapses on motor neurons as well as the interneurons responsible for recurrent and reciprocal inhibition, two critical inhibitory circuits that control motor output. At the postnatal ages in which SMA symptoms develop, calbindin^+^ Renshaw cells (responsible for recurrent inhibition) and FoxP2^+^ Ia inhibitory interneurons (responsible for reciprocal inhibition) are known to receive monosynaptic proprioceptive synaptic inputs (Bikoff et al., 2016; Jankowska and Roberts, 1972; Mentis et al., 2010; Mentis et al., 2006; Siembab et al., 2010; Worthy et al., 2023). Renshaw cells also receive additional, major excitatory drive through recurrent collaterals of motor axons exiting the spinal cord (Alvarez et al., 1999; Eccles et al., 1954; Lamotte d’Incamps and Ascher, 2008; Moore *et al*., 2015; Renshaw, 1946). Motor axon synapses on Renshaw cells are first established in embryo and proliferate postnatally (Alvarez *et al*., 2013; Siembab *et al*., 2010). We therefore first examined whether proprioceptive and motor axon inputs on these interneurons are affected during early postnatal development in an SMA mouse model.

To investigate proprioceptive-mediated neurotransmission on Renshaw cells during early SMA, we utilized whole-cell patch clamp intracellular recordings using the *ex vivo* spinal cord preparation at P3/4 in wild type (WT) and SMA pups of the SMN-Δ7 mouse model (Le et al., 2005). We targeted Renshaw cells in a “blind” manner focusing on the ventral area close to the exit of the ventral root, where most Renshaw cells are located (Geiman et al., 2000; Mentis *et al*., 2006). Renshaw cells were defined as ventral interneurons with robust monosynaptic responses to ventral root (motor axon) stimulation. Monosynaptic excitatory postsynaptic potentials (EPSPs) elicited by suprathreshold stimulation of the L1 ventral root, revealed no difference between WT and SMA Renshaw cells (Fig. 1A-C). In striking contrast, dorsal root L1-mediated EPSPs were markedly reduced in SMA Renshaw cells (Fig. 1D-F). Current-to-voltage relationship (Fig. 1G) revealed that SMA Renshaw cells exhibited increased input resistance, time constant and concomitant reduction in rheobase compared to their WT counterparts (Fig. 1H). The resting potential, voltage threshold and capacitance were not significantly different between the two genotypes (Fig. 1H). Neurobiotin (Nb) was injected intracellularly in Renshaw cells and visualized *post hoc* to confirm anatomically the identity of the recorded neurons as Renshaw cells. This approach, combined with immunohistochemistry against the vesicular acetylcholine transporter (VAChT) - a well-established marker for motor neuron axon collateral synapses - and the vesicular glutamate transporter one (VGluT1) - a protein present in proprioceptive synapses - validated morphologically the identity of the recorded neurons as Renshaw cells (Fig. 1I,J) and confirmed the presence of motor neuron axon collaterals and proprioceptive synapses on their somato-dendritic compartments.

**Fig. 1.**
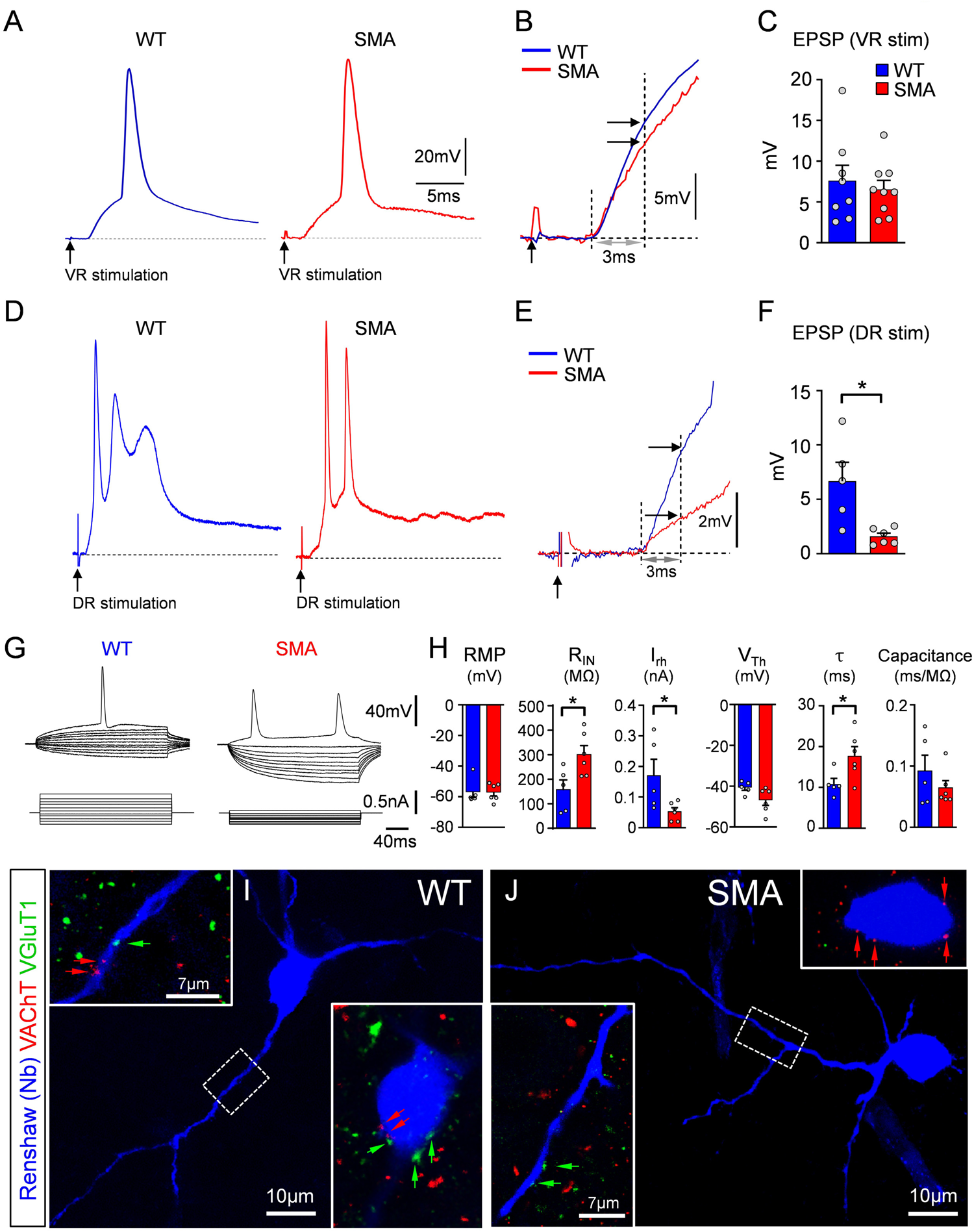
Reduced activation of Renshaw cells by proprioceptive synapses at the onset of SMA. **(A)** Excitatory postsynaptic potentials (EPSPs) in Renshaw cells following ventral root stimulation in wild type (blue) and SMA (red) mice. **(B)** Superimposed EPSPs from (A) at an expanded time scale. The maximum monosynaptically-induced EPSP amplitude was measured at 3ms from its onset (vertical dotted line and arrows). **(C)** Amplitude of VR-induced EPSPs between wild type (WT; n=8 Renshaws, N=8 mice) and SMA (n=9 Renshaw cells, N=9 mice) Renshaw cells (p=0.62, unpaired two-tailed t-test). **(D)** EPSPs in Renshaw cells following dorsal root (DR) stimulation in wild type (blue) and SMA (red) mice. **(E)** Superimposed EPSPs from (B) at an expanded time scale. Similar to (B), the EPSP amplitude was measured at 3ms from its onset. **(F)** Values of DR-induced EPSP amplitude in wild type (blue; n=5, N=5) and SMA (red; n=6, N=6) Renshaw cells. * p=0.0115, unpaired two-tailed t-test. Point of stimulation is denoted by a black arrow. **(G)** Superimposed traces of voltage responses (top traces) to current injections (bottom traces) in a wild type (WT) and a SMA Renshaw cell. **(H)** Resting membrane potential (RMP), input resistance (R_IN_), rheobase (I_rh_), voltage threshold (V_Th_), time constant (τ) and capacitance for wild type and SMA Renshaw cells. * p<0.05, all unpaired two-tailed t-tests (WT: n=5, N=5; SMA: n=6, N=6; p=0.0251 in R_IN_, p=0.0395 in I_rh_, and p=0.0376 in τ respectively). **(I,J)** Recorded cells intracellularly filled with Neurobiotin (visualized in blue) in wild type (I) and a SMA (J) mice together with immunoreactivity against VAChT+ (red) and VGluT1+ (green) synapses (respective arrows) in apposition onto their somata (insets) and dendrites (dash box and insets).

Together, these results indicate that proprioceptive neurotransmission on Renshaw cells is reduced by SMN deficiency at the onset of disease, like vulnerable SMA motor neurons. Moreover, this deficit is specific to the proprioceptive input, as excitation from motor neurons is unaffected at this early age. Lastly, some passive and active membrane properties in SMA Renshaw cells have been altered to signify a higher excitability state.

### Decrease in proprioceptive synapses on Renshaw cells at the onset of SMA

To investigate the extent of synaptic changes from motor neuron axon collaterals and proprioceptive neurons on Renshaw cells, we performed synaptic density measurements on Neurolucida reconstructed Renshaw neurons in WT and SMA mice at P3. Motor neuron axon collaterals were marked by retrograde fill with Cascade Blue dextran (Fig. 2A,D,I, and L; see Methods for details), while antibodies against VAChT validated these appositions as synapses on Renshaw cells. Renshaw cells were identified by their location in the ventral horn (yellow oval dotted area in Fig. 2A) and calbindin immunoreactivity (Alvarez *et al*., 1999; Carr et al., 1998). Using confocal microscopy (Fig. 2A,B,D,and E) and Neurolucida reconstructions of Renshaw cells (Fig. 2C,F), we found no significant difference in motor axon cholinergic synaptic coverage either on their soma (Fig. 2G) or their proximal dendrites (Fig. 2H) between WT and SMA mice at P3. In contrast, proprioceptive synapses marked by VGluT1 and parvalbumin (Fig. 2I-K, and L-N) were significantly reduced on both somatic (Fig. 2O) and proximal dendritic compartments (Fig. 2P) of Renshaw cells in SMA mice. Thus, SMA Renshaw cells receive fewer proprioceptive synapses, indicating that proprioceptive synaptic loss is not restricted to SMA motor neurons, but affects other spinal cord neurons.

**Fig. 2.**
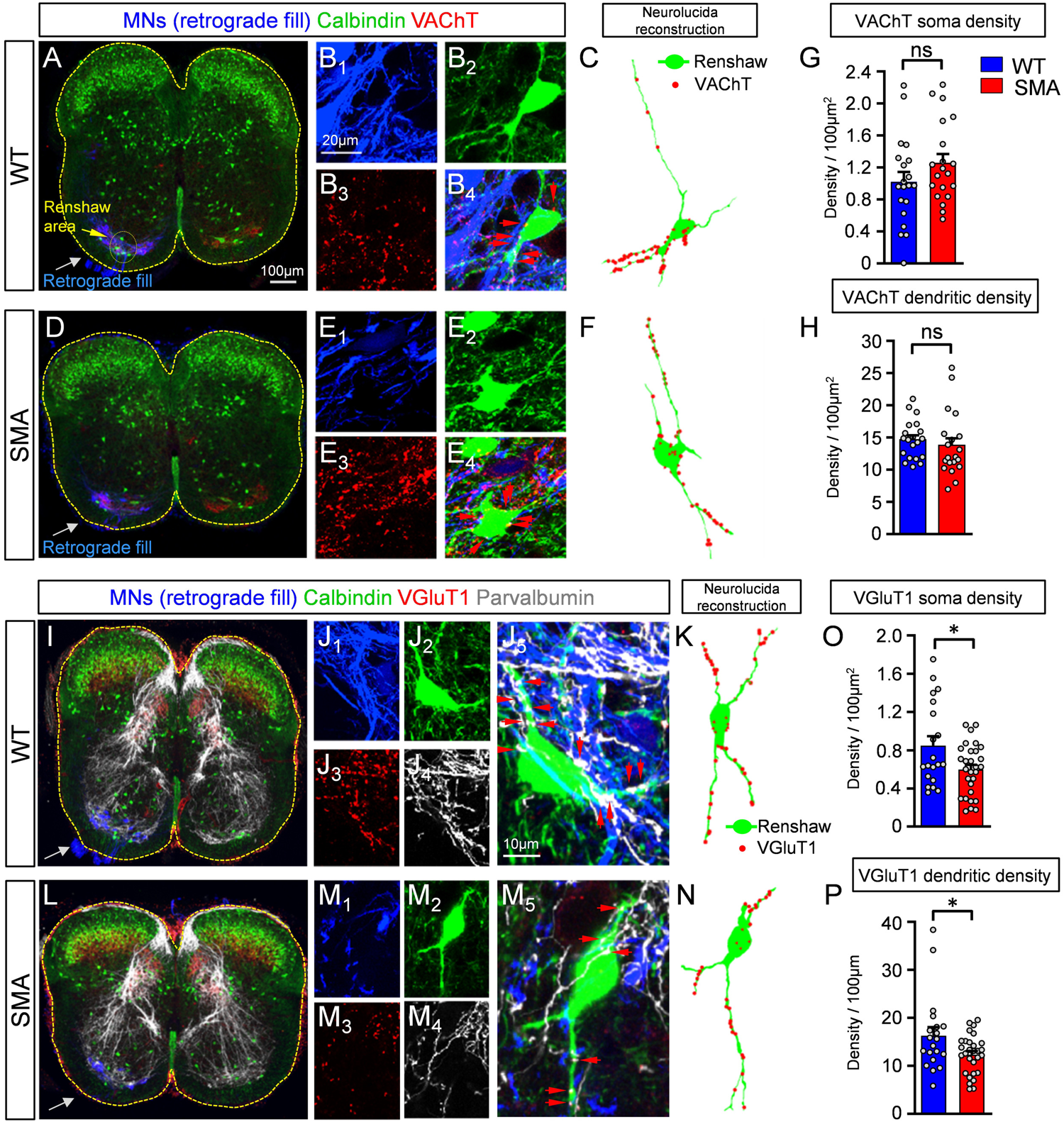
Decrease in synaptic density of proprioceptive synapses on Renshaw cells at the onset of SMA. **(A)** Low magnification image of a spinal cord showing immunoreactivity against calbindin (green), VAChT (red) and retrogradely filled motor neurons (in blue) in one side of the cord at P3. **(B_1-4_)** Higher magnification images of retrogradely filled motor neuron axon collaterals (B_1_, blue), a calbindin+ Renshaw cell (B_2_, green), VAChT immunoreactivity (B_3_, red) and a merged image (B_4_). **(C)** Neurolucida reconstruction of a wild type Renshaw cell (in green) with appositions from motor neuron axon collaterals (in red). **(D)** as in (A) but for an SMA mouse at P3. **(E_1-4_)** as in B_1-4_ for an SMA spinal cord. **(F)** Neurolucida reconstruction of a SMA Renshaw cell, similar to (C). **(G)** Density of cholinergic VAChT+ synapses on the somata of wild type and SMA Renshaw cells at P3. Each dot represents one Renshaw cell (n = 10 cells per animal and genotype). **(H)** Density of cholinergic VAChT+ synapses on the dendrites of wild type and SMA Renshaw cells at P3 (n as in G). There was no significant difference in H or G between the two groups (unpaired two-tailed t-test). **(I)** Low magnification image of a wild type spinal cord showing unilateral retrograde fill of motor neurons (blue), calbindin (green), VGluT1 (red) and parvalbumin (white) immunoreactivity. **(J_1-5_)** Higher magnification images of motor neuron axon collaterals (J_1_, blue), a calbindin+ Renshaw cell (J_2_, green), VGluT1 (J_3_, red), parvalbumin (J_4_, white) and their merged image (J_5_). Red arrows in J5 denote proprioceptive synapses (VGluT1+ and parvalbumin+) on the soma and dendrites of the Renshaw cell. **(K)** Neurolucida reconstruction of the Renshaw cell shown in J1-5 with VGluT1+/parvalbumin+ synaptic appositions (red). **(L)** Low magnification of a SMA spinal cord at P3, as in (I). **(M_1-5_)** Higher magnification images for a SMA Renshaw cell, as in (J_1-5_). **(N)** Neurolucida reconstruction of a SMA Renshaw with VGluT1+ synaptic appositions, as in (K). **(O)** VGluT1+/parvalbumin+ synaptic density on the somata of wild type and SMA Renshaw cells at P3; * p=0.0165, unpaired two-tailed t-test (n=19 WT N=2 mice and n=29 SMA N=3 mice). **(P)** VGluT1+/parvalbumin+ synaptic density on the dendrites of the same Renshaw cells; * p=0.0245, unpaired two-tailed t-test.

### Unexpected incursion of corticospinal VGluT1^+^ synapses on Renshaw cells at the end stage of SMA

The loss of proprioceptive synapses from motor neurons in SMA mice is progressive and follows synapse dysfunction (Fletcher et al, 2017). To investigate whether a similar progressive loss of proprioceptive synapses occurs on Renshaw cells, we examined VGluT1 synaptic coverage on Renshaw cells around vulnerable L1 SMA motor neurons at disease end stage (P11). To do so, a subset of vulnerable motor neurons was retrogradely labelled with CTb-488 from the iliopsoas muscle. Surprisingly, VGluT1^+^ synapses on SMA Renshaw cells were significantly increased on their soma and proximal dendrites at P11 (Fig. 3A-H). In the second postnatal week parvalbumin content diminishes in the central axons of proprioceptive afferents and upregulates in the axons of many spinal neurons (Siembab *et al*., 2010), preventing us from using parvalbumin to confirm the sensory origins of VGluT1+ synapses at P11. Thus, the unexpected increase in VGluT1^+^ synapse coverage could be due to either an increase of proprioceptive synaptic coverage at end stage of SMA, as previously suggested (Thirumalai et al., 2013), or due to a proliferation of VGluT1^+^ synapses from a different source.

**Fig. 3.**
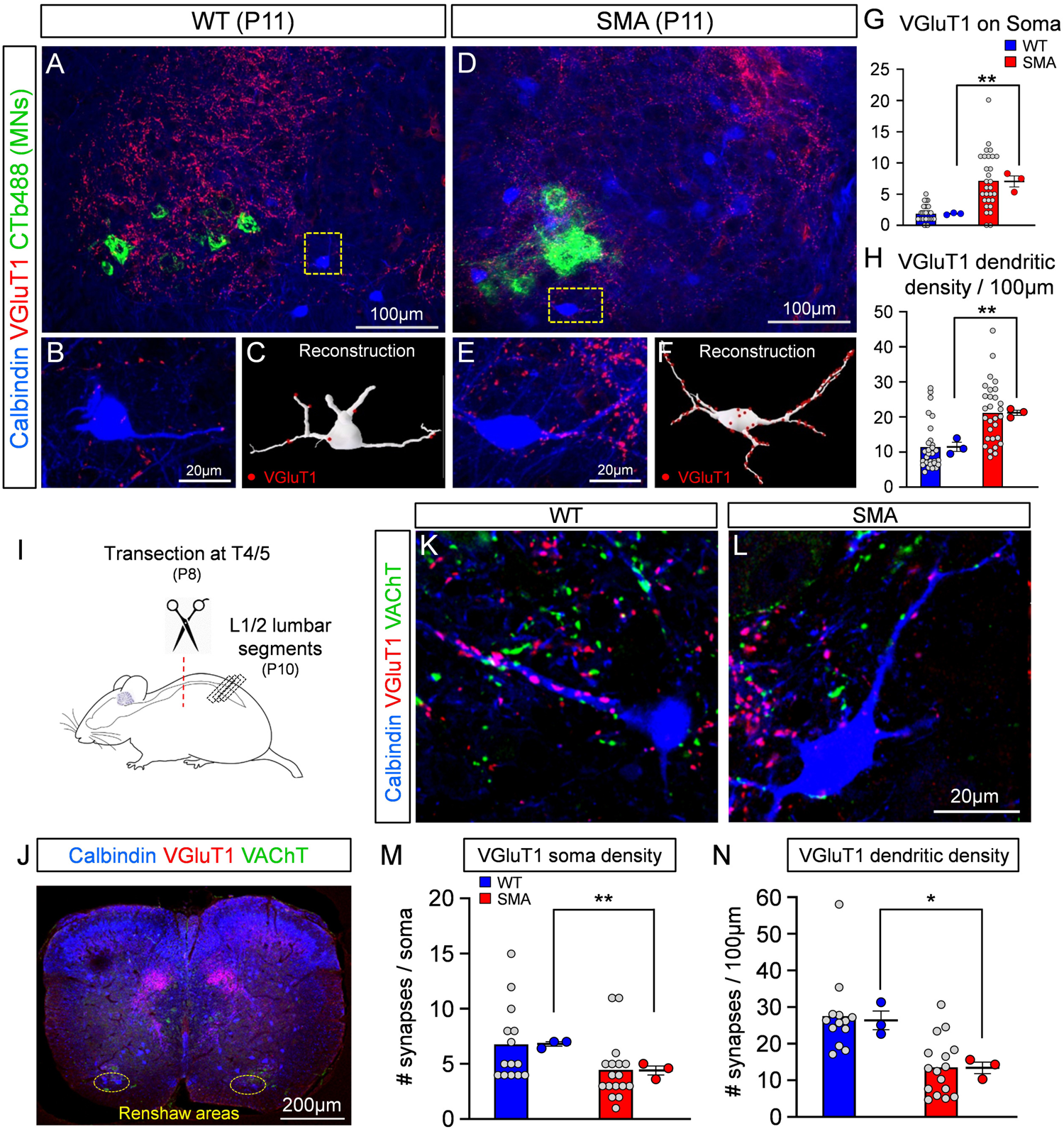
Unexpected increase of VGluT1+ synaptic coverage on Renshaw cells at the end stage of SMA. Low magnification images of the ventral horn showing iliopsoas motor neurons, labelled by retrograde muscle injection with CTb488 (green), calbindin (blue) and VGluT1 (red) immunoreactivity in a wild type **(A)** and a SMA **(D)** mouse, at P11. Single plane confocal images of a calbindin (blue) Renshaw cell and VGluT1+ synapses (red) in the wild type **(B)** and a SMA **(E)** mouse, from the dotted boxes in (A) and (D). Neurolucida reconstruction (**C**, white) of wild type Renshaw cell shown in (B), and a SMA Renshaw cell (**F**, white). VGluT1+ synaptic appositions are indicated as red dots. **(G)** VGluT1+ synaptic density on the somata of wild type (n=29 cells, N=3 mice) and SMA (n=29 cells, N=3 mice) Renshaw cells; ** p=0.0044, unpaired two-way t-test. **(H)** VGluT1+ synaptic density on the dendrites of wild type (n=29 cells, N=3 mice) and SMA (n=29, N=3 mice) Renshaw cells; ** p=0.0029, unpaired two-tailed t-test. Statistical comparison is between the average for each mouse in each genotype. **(I)** Experimental protocol for the spinal cord transection performed in the thoracic segment T4/5 at P8, followed by morphological examination of Renshaw cells located in the L1/2 spinal segments at P10. **(J)** Low magnification image of a transverse section of a spinal cord labelled with calbindin (blue), VGluT1 (red) and VAChT (green) antibodies. The Renshaw areas are shown bilaterally in the dotted oval circles. High magnification confocal images of a wild type **(K)** and a SMA **(L)** Renshaw cell (blue), together with VGluT1 (red) and VAChT (green) immunoreactivity after spinal transection. **(M)** VGluT1+ synaptic density on the somata of wild type (n=14, N=3) and SMA (n=17, N=3) Renshaw cells at P11; ** p=0.0065, unpaired two-tailed t-test. **(N)** VGluT1+ synaptic density on the dendrites of wild type (n=14, N=3) and SMA (n=17, N=3) Renshaw cells at P11; * p=0.0127, unpaired two-tailed t-test.

It has been previously established that VGluT1^+^ synapses on motor neurons are exclusively of proprioceptive origin (Alvarez et al., 2004; Hughes et al., 2004; Mentis et al., 2006; Rotterman et al., 2014). We have also reported that Renshaw cells received functional VGluT1^+^ synapses which are of proprioceptive origin in neonates (Mentis et al., 2006). However, adult mouse Renshaw cells also receive VGluT1^+^ synaptic contacts of corticospinal origin (D’Acunzo et al., 2014). Moreover, competition between VGluT1^+^ synapses originated in sensory afferents or corticospinal axons can remodel VGlut1^+^ synapse organization on spinal cord neurons (Jiang et al., 2016). To investigate whether early deficits in VGluT1^+^ proprioceptive synapses over developing Renshaw cells result in increased VGluT1^+^ corticospinal synapses, we injected an AAV9-GFP vector bilaterally in the cortex of newborn (P0) mice (Suppl. Fig. 1A_1_). We then examined morphologically its presence on Renshaw cells in the L1/2 lumbar segments from WT and SMA mice at P10. After verification of successful injection of AAV9-GFP in the cortex (Suppl. Fig. 1A_2_), we determined that many Renshaw cells received GFP+ and VGluT1+ synapses on soma and dendrites in both WT and SMA spinal cords (Suppl. Fig. 1B,C). This indicates that Renshaw cells receive VGluT1^+^ corticospinal synapses at P10.

To quantify the extent of proprioceptive-derived VGluT1^+^ synapses on SMA Renshaw cells, we performed spinal cord transections at the 4^th^/5^th^ thoracic segment in WT and SMA mice at P8 and examined the number of VGluT1^+^ synapses remaining after elimination of corticospinal synapses on Renshaw cells at P10 (Fig. 3I,J). The success of the bilateral spinal cord transection was verified by histological examination of lesion completeness and verification of the spinal segment transected at P10. After removal of synapses from descending systems, we found that SMA Renshaw cells received significantly fewer VGluT1 synapses compared to similarly injured WT mice at P10 (Fig. 3K,L) on the soma (Fig. 3M) and more significantly, on proximal dendrites (Fig. 3N). We interpret these remaining VGluT1^+^ synapses and their depletion as specifically proprioceptive. The observed depletion is also opposite to the expected plasticity of VGluT1^+^ proprioceptive synapses on spinal interneurons of WT animals after removal of the corticospinal tract, which should increase in density based on previous studies (Goltash et al., 2023; Hollis et al., 2015). Thus, removing VGluT1^+^ synapses from proprioceptors on Renshaw cells in SMA mice also interferes with their expected plasticity (increase) after corticospinal tract injury.

Cholinergic VAChT+ synaptic densities on dendrites of Renshaw cells were similar in WT and SMA mice, and unaffected by spinal cord transection (Suppl. Fig. 1D-F). However, VAChT^+^ synapses on the cell body of Renshaw cells were significantly increased in SMA mice. This is likely due to the reported synaptic competition during early development between VGluT1^+^ proprioceptive synapses and motor axon VAChT^+^ synapses on proximal somatodendritic regions of Renshaw cells (Siembab et al., 2016).

Taken together, these results indicate that proprioceptive synaptic coverage on Renshaw cells is reduced in SMA mice both at the onset and end stage of disease. Additionally, the loss of excitatory proprioceptive synapses in SMA appears to enact a maladaptive increase of corticospinal and possibly motor axon synapses, which now occupy synaptic space, made available by the loss of proprioceptive input.

### Loss of proprioceptive synapses in SMA renders putative Ia inhibitory interneurons hyperexcitable

Ia inhibitory interneurons are responsible for reciprocal inhibition of antagonistic motor pools and provide an equally powerful source of inhibition to the cell body and proximal dendrites of motor neurons (Hultborn et al., 191; Jankowska and Roberts, 1972; Jankowska, 1992). Ia inhibitory interneurons originate from V1 and V2b spinal interneurons genetic classes (Alvarez et al., 2005; Zhang et al., 2014) and those derived from V1s can be characterized by expression of the transcription factor Foxp2 (Benito-Gonzalez and Alvarez, 2012; Worthy *et al*., 2023). As expected, since V1 interneurons also include Renshaw cells, ∼80% of the inhibitory synapses on adult mouse iliopsoas motor neuron cell bodies originate from combined V1 and V2b interneurons (Zhang et al., 2014). Of those synapses, ∼65% originate from Renshaw cells and other V1 interneurons, including V1-derived Ia inhibitory interneurons, whereas synapses from V2b interneurons represent less than 25% (Worthy *et al*., 2023; Zhang *et al*., 2014). Ia inhibitory interneurons are a large population of spinal interneurons and receive strong monosynaptic Ia afferent proprioceptive input (Hultborn et al., 1971; Hultborn and Udo, 1972; Siembab *et al*., 2010). We therefore investigated ventral horn interneurons - other than Renshaw cells - with monosynaptic responses to dorsal root stimulation (which is of proprioceptive origin if neurons are located in the ventral horn) to analyse whether they are affected in SMA mice. We recorded intracellularly from ventral interneurons using the *ex vivo* spinal cord from WT and SMA mice at P3/4. These neurons were identified as putative Ia interneurons based on: i) location just adjacent to motor pools, an area previously reported to be enriched with Ia interneurons (Benito-Gonzalez and Alvarez, 2012; Jankowska and Lindström, 1972; Worthy et al., 2023), ii) a monosynaptic response following stimulation of the homosegmental dorsal root (L1 or L2), and iii) absence of monosynaptically-mediated EPSP responses following ventral root stimulation. Based on these criteria, recordings from three WT and three SMA putative Ia inhibitory interneurons revealed that the SMA interneurons exhibited signs of hyperexcitability (Fig. 4A) as shown by the significant increase in input resistance (R_n_; Fig. 4B) and reduction in rheobase (I_Rh_; Fig. 4C), while the voltage threshold (V_Thr_) was unchanged between WT and SMA mice (Fig. 4D). Importantly, the monosynaptic EPSPs after sensory fiber stimulation were significantly reduced in SMA Ia neurons (Fig. 4E), an observation akin to that detected in SMA Renshaw cells (Fig. 1F). Notably, spinal interneurons that could not be activated monosynaptically by either proprioceptive fibers or motor neuron axon collaterals, did not reveal any significant differences in either passive or active intrinsic properties between WT and SMA mice (Suppl. Fig. 2A,B).

**Fig. 4.**
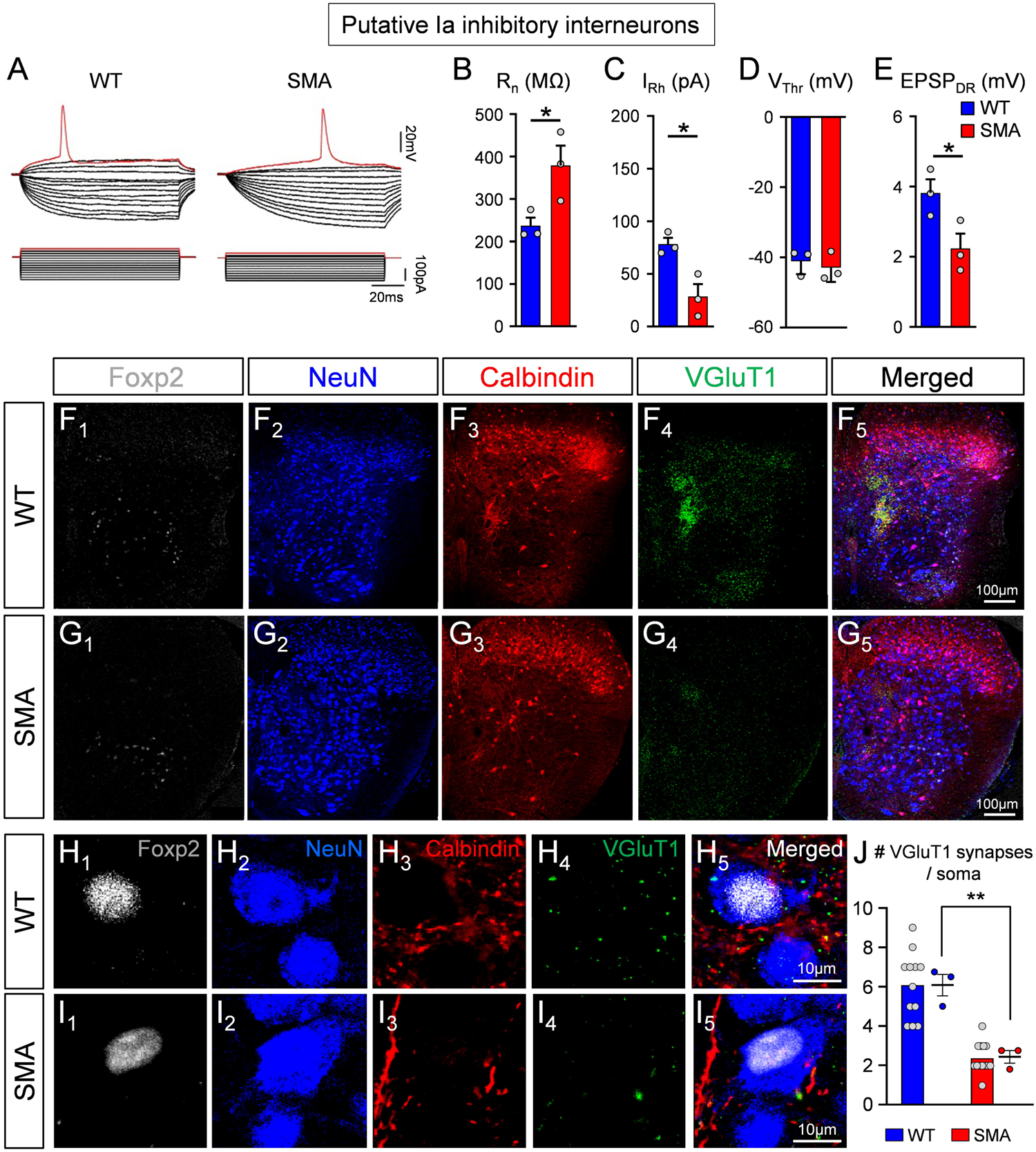
Putative Ia inhibitory interneurons are hyperexcitable and receive fewer proprioceptive synapses in SMA at the disease onset. (A) Superimposed voltage responses (top traces) to current injections (bottom traces) for a putative wild type (left) and a SMA (right) Ia inhibitory interneuron. The input resistance (B), rheobase (C), voltage threshold (D) and EPSP amplitude (E) in wild type (blue; n=3, N=3) and SMA (red, n=3; N=3) putative Ia inhibitory interneurons were significantly different; * p=0.048 in (B), p=0.019 in (C), p=0.049 in (E), unpaired two-tailed t-test. (F, G) Low magnification images from one side of the spinal cord of a wild type (F_1-5_) and a SMA (G_1-5_) mouse, showing Foxp2 (white, F_1_, G_1_), NeuN (blue, F_2_, G_2_), calbindin (red, F_3_, G_3_), VGluT1 (green, F_4_, G_4_) immunoreactivity, as well as the merged image (F_5_, G_5_). (H, I) Higher magnification of Ia inhibitory interneurons, showing calbindin+ and VGluT1+ synapses in a wild type (H_1-5_) and a SMA (I_1-5_) mouse. (J) The number of VGluT1+ synapses on the soma of Ia inhibitory interneurons in wild type (n=12 cells, N=3 mice) and SMA (n=13 cells, N=3 mice) mice differ at P4; *** p=0.004, unpaired two-tailed t-test. Statistical comparison performed between WT and SMA mice.

The above results were complemented by anatomical analyses of proprioceptive-derived VGluT1^+^ synapses on histologically identified Ia inhibitory interneurons at the onset of SMA (P3). We performed a synaptic density analysis on NeuN demarcated somata of spinal interneurons that fulfilled the following criteria: i) nuclear presence of Foxp2 (Benito-Gonzalez and Alvarez, 2012); and ii) receive convergent calbindin^+^ and VGluT1^+^ synapses from Renshaw cells and muscle proprioceptors (Fig. 4F-I), respectively. These Ia inhibitory interneurons received significantly fewer synapses in P3 SMA mice compared to their WT counterparts (Fig. 4J).

Taken together, these results suggest that proprioceptive neurotransmission is reduced on spinal interneurons in parallel to motor neurons at the onset of SMA resulting in similar downstream alterations in the excitability of their postsynaptic neuronal targets.

### Inhibitory synaptic strength is higher in SMA motor neurons

We next investigated whether Renshaw cells in SMA have altered their firing ability. Following current injection, we determined that SMA Renshaw cells exhibit a higher frequency in repetitive firing (Fig. 5A,B). To address whether the hyperexcitable Renshaw cells in SMA mice have higher activity, we quantified the spontaneous firing frequency of Renshaw cells at their own resting potential in WT and SMA mice at P3/4 (Fig. 5C) and found that SMA Renshaw cells exhibited a significantly increase in spontaneous firing (Fig. 5D). These results indicate that SMA motor neurons may receive increased inhibitory drive. The increased excitability of various classes of inhibitory interneurons presynaptic to motor neurons suggest that SMN deficient motor neurons may be modulated by overactive interneurons that mature inhibitory synapses of higher strengths. To directly test this possibility, we recorded miniature inhibitory postsynaptic currents (mIPSCs) in L1 or L2 vulnerable SMA motor neurons in both WT and SMA mice at P3/4 using the *ex vivo* spinal cord preparation. mIPSCs of GABAergic and/or glycinergic origin were pharmacologically isolated with 1μM TTX, 10μM CNQX and 100μM APV, as well as 50μM mecamylamine, 50μM dHβE and 30μM D-tubocurarine, as previously reported (González-Forero and Alvarez, 2005). The glycinergic and/or GABAergic nature of recorded mIPSCs was verified by the addition of 10μM bicuculline and 0.25μM strychnine (Suppl. Fig. 3A,B). SMA motor neurons exhibited a significantly greater amplitude and frequency of mIPSCs (Fig. 5E,F,G). These results demonstrate that vulnerable SMA motor neurons receive inputs from inhibitory synapses of higher strength compared to WT motor neurons at the onset of the disease.

**Fig. 5.**
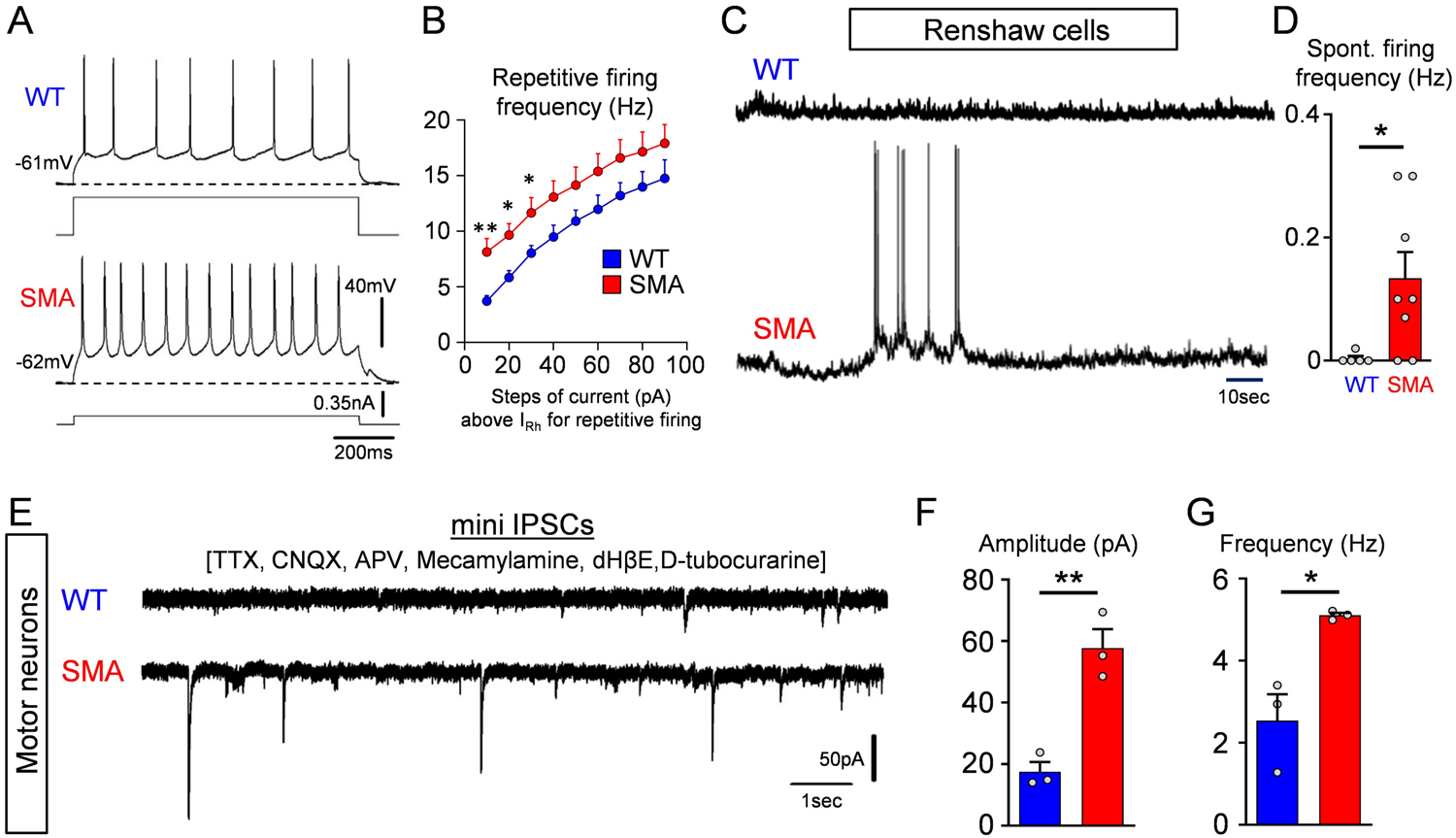
SMA motor neurons receive increased inhibitory synaptic drive. **(A)** Repetitive firing following current injection at 40 pA above threshold in a wild type and a SMA Renshaw cell at P4. **(B)** Average firing frequency after current injection in wild type (n=5 cells, N=5 mice) and SMA (n=5 cells, N=5 mice) Renshaw cells at P4. ** p=0.009 (10pA step), * p=0.013 (20pA step), * p=0.045 (30pA step), unpaired two-tailed t-test. **(C)** Spontaneous voltage activity in a wild type and a SMA Renshaw cell at P4. **(D)** Spontaneous firing frequency in wild type and SMA Renshaw cells. * p=0.0377, unpaired two-tailed t-test (WT: n=5 cells, N=5 mice; SMA: n=8 cells, N=8 mice). **(E)** Pharmacologically isolated mIPSCs in a wild type and SMA motor neuron located from the L2 spinal segment at P4. Amplitude **(F)** and frequency **(G)** of mIPSCs in wild type (n=3 cells, N=3 mice) and SMA (n=3 cells, N=3 mice) L2 motor neurons at P4. ** p=0.004, * p=0.016, unpaired two-tailed t-test.

### Increased incidence of glycinergic and GABAergic synapses on vulnerable SMA motor neurons

To investigate possible changes in synaptic coverage by inhibitory inputs we examined glycinergic and GABAergic synapses on L1/2 motor neurons in WT and SMA mice at P4. Presynaptic GABAergic boutons were identified by GAD65/67 antibodies, whereas glycinergic synapses were marked by GlyT2 antibodies (Fig. 6A-B and 6D-D, respectively). Both types of synapses are associated with postsynaptic gephyrin (Colin et al., 1998; Geiman et al., 2002; Todd et al., 1995). Gephyrin organizes the postsynaptic receptor clusters and perfectly matches the presynaptic active zone where the inhibitory neurotransmitter is released (Alvarez, 2017). We found a significant increase in the number of GAD65/67^+^ contacts around somata of L1/2 motor neurons in SMA compared to WT controls and in the number of inhibitory synaptic sites, marked by gephyrin, that are associated with presynaptic GAD65/67 (Fig. 6G-H). The number of GAD65/67 synaptic sites revealed by gephyrin increased more than the number of GAD65/67 boutons because more boutons became associated with more than one synaptic complex. Similarly, the number of glycinergic bouton appositions (GlyT2^+^) and gephyrin synaptic sites opposite to GlyT2^+^ boutons significantly increased in SMA compared to WT controls (Fig. 6I-J). Moreover, individual synaptic boutons showed increased frequency of multiple release sites (marked by independent gephyrin clusters) in SMA motor neurons compared to WTs.

**Fig. 6.**
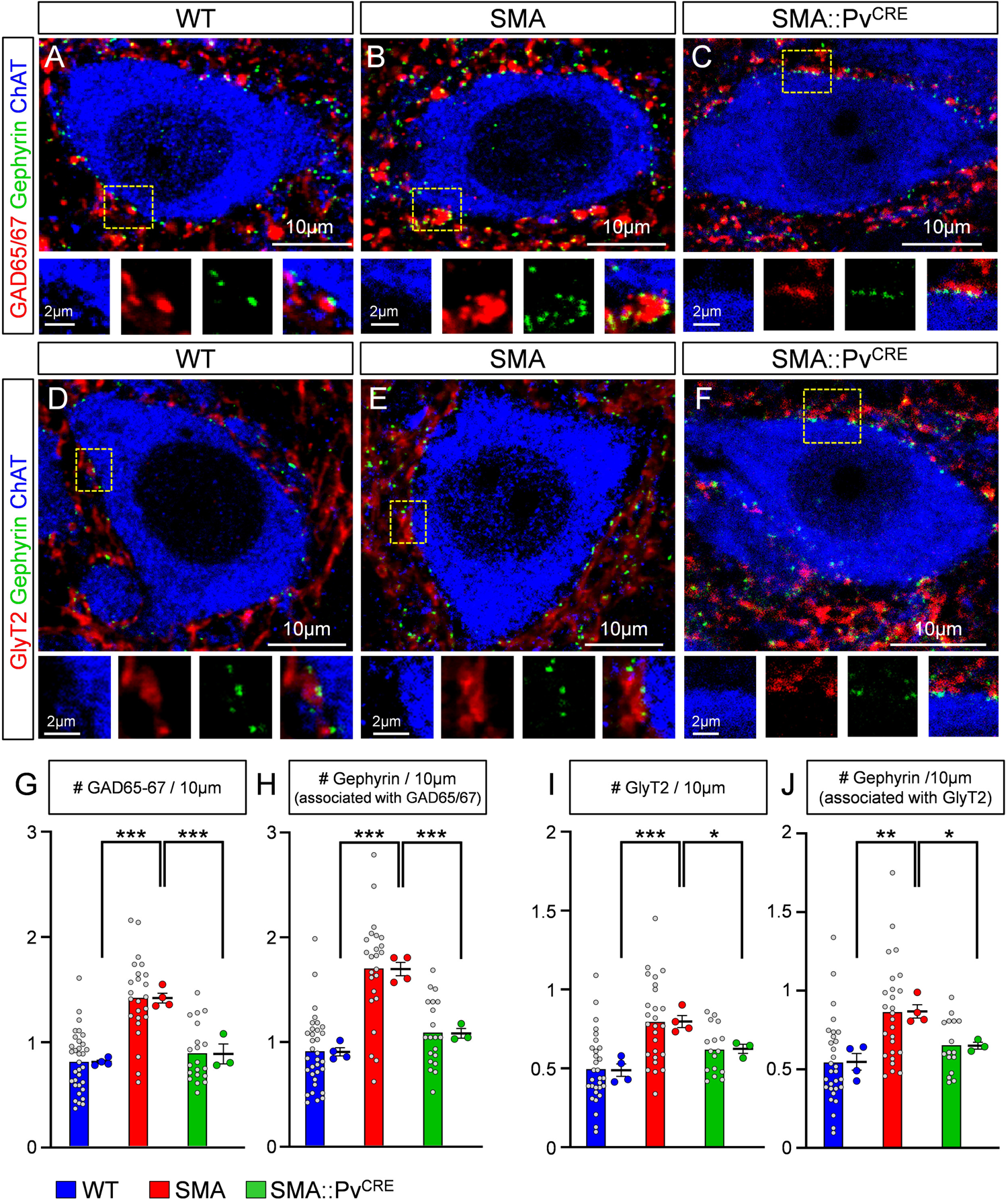
SMA motor neurons are covered by a higher number of GABAergic and glycinergic synapses. **(A-C)** Single plane confocal images from a wild type (A), a SMA (B) and SMA::Pv^CRE^ (C) motor neuron showing GAD65/67 (red), gephyrin (green) and ChAT (blue) immunoreactivity at P4. Insets at the bottom are higher magnification images from the dotted boxed area, showing the individual fluorochromes. **(D-F)** Single plane confocal images from a wild type (D), a SMA (E) and a SMA::Pv^CRE^ (F) motor neuron showing GlyT2 (red), gephyrin (green) and ChAT (blue) immunoreactivity at P4. Insets at the bottom are higher magnification images from the dotted boxed area, showing the individual fluorochromes. **(G)** The number of GAD65/67 synapses per 10 μm of motor neuron (MN) membrane in wild type (n=33 MNs, N=4 mice), SMA (n=24 MNs, N=4 mice) and SMA::Pv^CRE^ (n=20 MNs, N=3 mice) motor neurons. *** p<0.0001 WT vs SMA, *** P=0.0004 SMA vs SMA::Pv^CRE^, OneWay ANOVA, Tukey’s *post hoc* test. **(H)** The number of gephyrin clusters associated with GAD65/67+ synapses per 10μm of motor neuron membrane in wild type and SMA motor neurons (“n” and “N” identical as in G). *** p<0.0001 WT vs SMA, *** P=0.0001 SMA vs SMA::Pv^CRE^, OneWay ANOVA, Tukey’s *post hoc* test. **(I)** The number of GlyT2 synapses per 10μm of motor neuron membrane in wild type (n=30 MNs, N=4 mice), SMA (n=26 MNs, N=4 mice) and SMA::Pv^CRE^ (n=16 MNs, N=3 mice) motor neurons. *** p=0.0008 WT vs SMA, * p=0.032 SMA vs SMA::Pv^CRE^, OneWay ANOVA, Tukey’s *post hoc* test. **(J)** The number of gephyrin clusters associated with GlyT2+ synapses per 10μm of motor neuron membrane in wild type and SMA motor neurons (“n” and “N” identical as in I). ** p=0.0018 WT vs SMA, * p=0.023 SMA vs SMA::Pv^CRE^, OneWay ANOVA, Tukey’s *post hoc* test. Statistical comparison was performed between the average values from mice.

Multiple release sites are a common feature of inhibitory synapses on adult motor neurons (Alvarez et al., 1997) and are usually interpreted as augmenting probability of release from single boutons (Alvarez, 2017). These results suggest more rapid maturation of inhibitory synapses with multiple release sites during early postnatal development on vulnerable motor neurons in SMA.

To investigate whether these changes in the number of inhibitory synapses originate from the reduction of proprioceptive synapses or occurs independently, we quantified GABAergic and glycinergic synapses in SMA::Pv^CRE^ mice in which SMN was selectively restored in proprioceptive neurons by genetic means using the SMA Conditional Inversion mice (Lutz et al., 2011) crossed with Pv-Cre mice, that we validated in a previous report (Fletcher et al., 2017). Importantly, we found that both GABAergic (GAD65/67) (Fig. 6A-C) and glycinergic (GlyT2) (Fig. 6D-F) synapses were rescued to levels similar to those of WT mice (Fig. 6G-J). This demonstrates that the aberrant increase of inhibitory synapse inputs is a consequence of the loss of excitatory proprioceptive synapses.

### Downregulation of gephyrin in motor neurons *in vivo* rescues cellular phenotypes and provides behavioural benefit in SMA mice

In parallel with the reduced excitatory drive from proprioceptive neurons, excessive inhibition could further impair recruitment of motor neurons and contribute to muscle weakness and/or paralysis in SMA mice. To test this possibility, we investigated whether reducing inhibitory drive on SMA motor neurons improves the disease phenotype. To alleviate the inhibitory drive selectively on motor neurons, we developed an AAV9-based strategy for shRNA-mediated knockdown of gephyrin *in vivo* (Suppl. Fig. 4A). We have previously shown that intracerebroventricular (i.c.v.) injection of AAV9 at P0 transduces motor neurons and a few glial cells but no interneurons in the spinal cord (Simon et al., 2017). Accordingly, we quantified the percentage of motor neurons transduced by the AAV9-Geph_RNAi_ vector which also expresses GFP, following i.c.v. injection at P0(Suppl. Fig. 4B_1-3_) and found that 65-70% WT motor neurons and 60-65% SMA motor neurons were transduced when examined at P11 (Suppl. Fig. 4C). We then tested the effects of AAV9-Geph_RNAi_ injection on gephyrin expression in L1/2 motor neurons using immunohistochemistry at both P4 (Fig. 7A_1-4_) and P11 (Suppl. Fig. 4D). We found that the number of gephyrin clusters in motor neuron somata was significantly reduced by ∼51% at P4 (Fig. 7B) and by ∼52% at P11 (Suppl. Fig. 4E) in WT motor neurons, similar to SMA motor neurons (∼54% at P4 and ∼75% at P11). Importantly, gephyrin knockdown reversed the increase in amplitude and frequency of mIPSCs in L1/2 motor neurons in SMA mice at P4 (Fig. 7C,D). These results demonstrate that knockdown of gephyrin in SMA motor neurons rescues the unwarranted increase in amplitude and frequency of the mIPSCs observed in SMA mice.

**Fig. 7.**
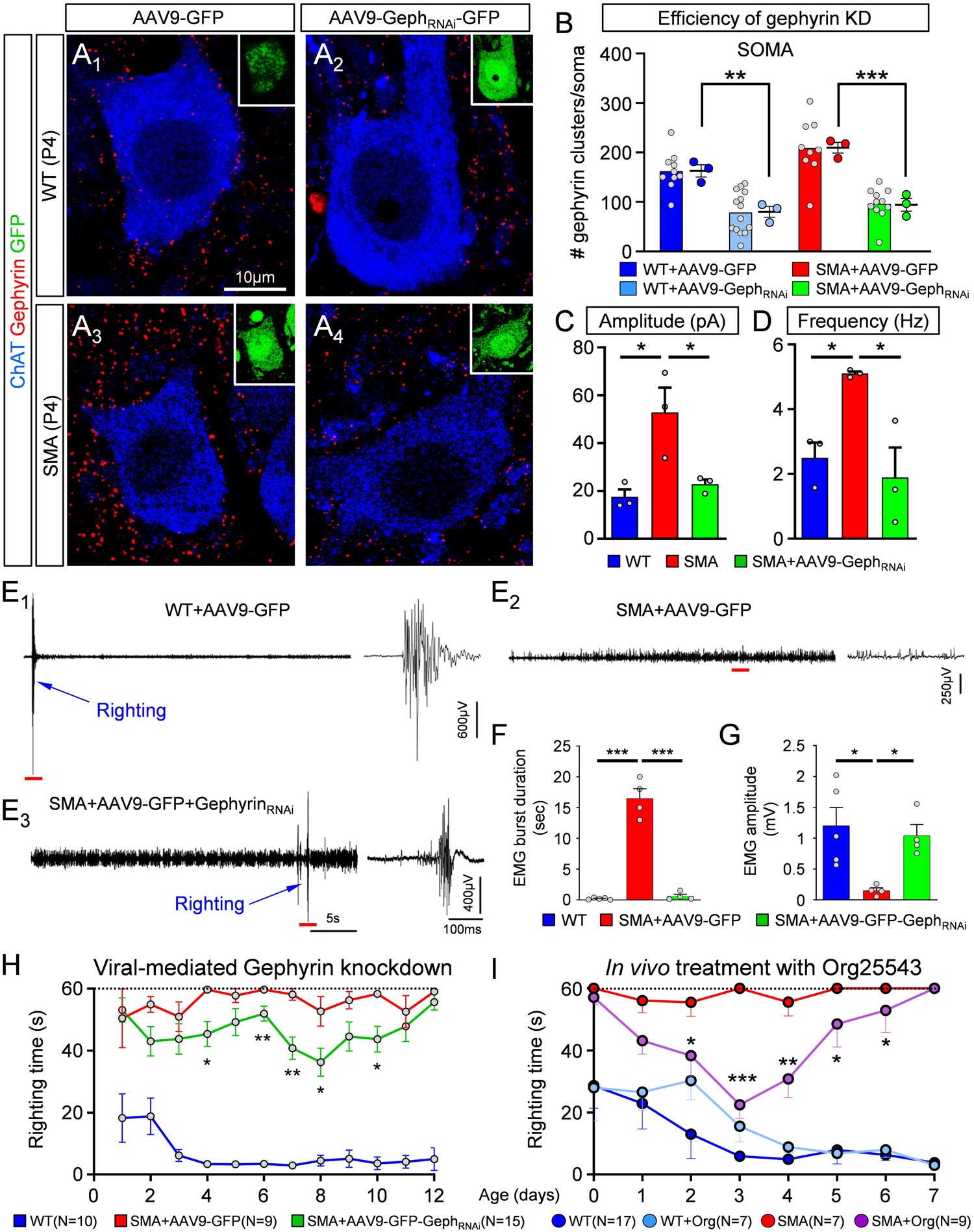
Knockdown of gephyrin *in vivo* abolishes the enhancement of inhibitory synapses in SMA motor neurons and provides phenotypic benefit in SMA mice. (A_1-4_) Single optical plane confocal images of wild type (A_1,2_) and SMA (A_3,4_) motor neurons transduced either with AAV9-GFP (A_1,3_) or with AAV9-Gephy_RNAi_-GFP (A_2,4_). ChAT (blue) and Gephyrin (red) immunoreactivity is shown for each case. Insets show GFP (green) expression in these motor neurons. Number of gephyrin clusters per motor neuron soma **(B)** for the four experimental groups shown in (A). WT+AAV9-GFP (n=10 MNs; N=3 mice), WT+AAV9-Geph_RNAi_-GFP (n=14 MNs; n=3 mice), SMA+AAV9-GFP (n=9 MNs, N=3 mice), SMA+AAV9-Geph_RNAi_-GFP (n=10 MNs, N=3 mice). ** p=0.0049, WT vs WT+AAV9-Geph_RNAi_; *** p=0.0006, SMA vs SMA+AAV9-Geph_RNAi_; OneWay ANOVA, Tukey’s multiple *post hoc* tests. Amplitude **(C)** and frequency **(D)** of mIPSCs recorded from L1/2 motor neurons in wild type (blue, n=3 MNs, N=3 mice), SMA (red, n=3 MNs, N=3 mice) and SMA+AAV9-Geph_RNAi_ (green, n=3 MNs, N=3 mice) mice at P3/4. In (**C**): * p=0.018, WT vs. SMA; * p=0.035, SMA vs. SMA+AAV9-Geph_RNAi_; OneWay ANOVA, Tukey’s *post hoc* test. In (**D**): *\** p=0.049, WT v SMA; * p=0.021, SMA v SMA+AAV9-Geph_RNAi_; OneWay ANOVA, Tukey’s *post hoc* test. (E_1-3_) *in vivo* EMG recordings from iliopsoas muscle during righting reflex in a control wild type (WT+AAV9-GFP), an SMA mouse treated with AAV9-GFP (SMA+AAV9-GFP) and an SMA mouse treated with AAV9-Geph_RNAi_-GFP at P10. Traces in the right-hand side are time-expanded samples taken from the long trace, denoted by the red bar. Righting of the mouse is denoted by the blue arrow. **(F)** Duration of EMG burst activity during the righting reflex test. *** p<0.0001 for both, WT+AAV9-GFP vs SMA+AAV9-GFP as well as SMA+AAV9-GFP vs SMA+AAV9-Geph_RNAi_-GFP; OneWay ANOVA, Tukey’s *post hoc* test. **(G)** Amplitude of EMG response during righting reflex test. * p=0.0167, WT+AAV9-GFP vs SMA+AAV9-GFP mice; * p=0.0500, SMA+AAV9-GFP vs SMA+AAV9-Geph_RNAi_-GFP mice; OneWay ANOVA, Tukey’s *post hoc* test. (H) Righting reflex times for control (WT+AAV9-GFP) mice (blue; N=10 mice), SMA mice treated with AAV9-GFP (red, N=9 mice) and SMA mice treated with AAV9-Geph_RNAi_-GFP (green, N=15 mice). * p<0.05, ** p<0.01, unpaired two-tailed t-tests for individual postnatal ages, for SMA+AAV9-GFP vs SMA+AAV9-Geph_RNAi_-GFP mice. **(I)** Righting reflex times following pharmacological *in vivo* treatment with Org25543. * p<0.05, ** p<0.01 and *** p<0.001, unpaired two-tailed t-tests between SMA and SMA+Org25543 mice for individual postnatal ages. (WT: N=17 mice; SMA: N=7 mice; WT+Org: N=7 mice; SMA+Org: N=9 mice).

Since Na-K-2Cl and K-Cl cotransporters are important regulators of intracellular chloride in neurons and determine the strength GABA or glycine neurotransmission (Kahle et al., 2010; Payne et al., 2003; Rivera et al., 1999), we investigated potential changes in NKCC1 and KCC2 expression in vulnerable (medial L5 motor neurons) and resistant (lateral L5 motor neurons) motor neurons in WT and SMA mice. Through a laser capture microdissection approach, as we previously reported (Simon *et al*., 2017), we collected at P4 the cytoplasm from WT and SMA motor neurons, labelled with CTb-488 retrogradely by *in vivo* intramuscular injection at birth. We then performed mRNA expression analysis by RT-qPCR and found no difference in either *Nkcc1* or *Kcc2* mRNAs between WT and SMA mice (Suppl. Fig. 4F,G). This suggests that the basic mechanisms of neuronal chloride homeostasis and inhibitory synapse driving forces are not altered between WT and vulnerable SMA motor neurons.

Next, we wanted to investigate whether lessening the inhibitory drive on vulnerable motor neurons by virally-mediated knockdown of gephyrin, would improve muscle function. To test this, we implanted a bipolar electrode in the iliopsoas/quadratus lumborum (IL/QL) muscles of WT and SMA mice and recorded electromyogram (EMG) activity (Fig. 7E). The IL/QL muscles are clinically relevant muscles innervated by vulnerable L1-L3 motor neurons and involved in righting ability (Fletcher *et al*., 2017; Mentis *et al*., 2011). At P11, WT, but not SMA, mice exhibited a rapid ability to right themselves within a couple of seconds, which was associated with short duration and high amplitude EMG activity from the IL/QL muscles (Fig. 7E_1_). In contrast, SMA mice exhibited low amplitude, continuous activity during the 60sec period of the test in which the pup was unable to right (Fig. 7E_2_). Knockdown of gephyrin resulted in a delayed but successful ability of SMA mice to right themselves, which was evident in the EMG recording that consisted of similar short bursts as recorded in WT mice but with significant delay from the start of the test (Fig. 7E_3_). On average, gephyrin knockdown resulted in a significantly shorter duration (Fig. 7F) and a higher amplitude of EMG activity (Fig. 7G) from the IL/QL muscles during the righting reflex.

The changes in EMG activity were associated to the behavioural phenotype of SMA mice. SMA mice injected with AAV9-Geph_RNAi_-GFP exhibited a modest but significant improvement in righting time as early as P4 compared to SMA mice treated with control AAV9-GFP (Fig. 7H). To test whether targeting inhibition pharmacologically could also provide behavioural benefit, we injected WT and SMA mice i.p. with 1.5mg/kg Org-25543, a GlyT2 inhibitor that crosses the blood brain barrier (Mingorance-Le Meur et al., 2013) and reduces glycinergic transmission (Al-Khrasani et al., 2019; Rousseau et al., 2008). The injections were performed daily, starting at birth until P7. Interestingly, we found a significant yet transient improvement in the righting ability of SMA mice treated with Org25543 (Fig. 7I). This treatment also resulted in robust improvement of weight gain (Suppl. Fig. 4I). Although interfering with gephyrin expression resulted in no benefit in the lifespan of SMA mice (Suppl. Fig. 4H), this can be explained by lethality in this animal model by additional mechanisms (Bevan et al., 2010; Shababi et al., 2012).

Collectively, these results indicate that excess inhibitory drive on spinal motor neurons contributes to motor dysfunction in SMA mice and provide proof-of-concept that targeting this deficits either genetically or pharmacologically has the potential to mitigate the disease phenotype.

## DISCUSSION

The mechanisms by which dysfunctional synaptic circuits result in neuropathology are poorly understood for many neurodegenerative diseases. SMA is a neurodegenerative disease characterized by early pathophysiology in sensory-motor circuits, which reduces excitatory drive to affected motor neurons (Mentis et al., 2011; Fletcher et al., 2017). Under normal synaptic homeostatic mechanisms, it is expected that the inhibitory synaptic strength be reduced in parallel to maintain motor neuron activity within narrow windows around predetermined firing set points. Failure to do so would further decrease the capacity of spinal motor circuits to recruit motor neurons and drive muscle activity, resulting in paralysed muscles. Here, we shed light on the interplay between excitation and inhibition in SMA motor neurons and their active participation in disease pathogenesis. The inhibitory synaptic drive on motor neurons is dominated by two principal inhibitory interneurons: Renshaw cells and Ia inhibitory interneurons (Worthy *et al*., 2023). We determined that premotor inhibitory spinal interneurons receive reduced excitatory drive from Ia afferents in SMA mice (Fig. 8). This resulted in increased excitability and also changes in their synaptic inputs, as shown for Renshaw cells receiving larger drive from descending systems (Fig. 8). Moreover, the expected homeostatically-mediated reduction in motor neuron inhibition by premotor inhibitory interneurons did not occur. Instead, vulnerable motor neurons received aberrantly increased inhibitory synaptic activity (Fig. 8). Importantly, genetic alleviation of the undue inhibition imposed on SMA motor neurons resulted in significant yet transient improvement in cellular and neuronal circuit function as well as behavioural benefits. Thus, our study identifies lack of excitation-inhibition (E/I) homeostasis as a major maladaptive event in SMA and suggests that the combined effect of reduced excitation and increased inhibition cooperatively diminish the capacity of premotor commands to recruit motor neurons and elicit muscle contractions.

**Fig. 8.**
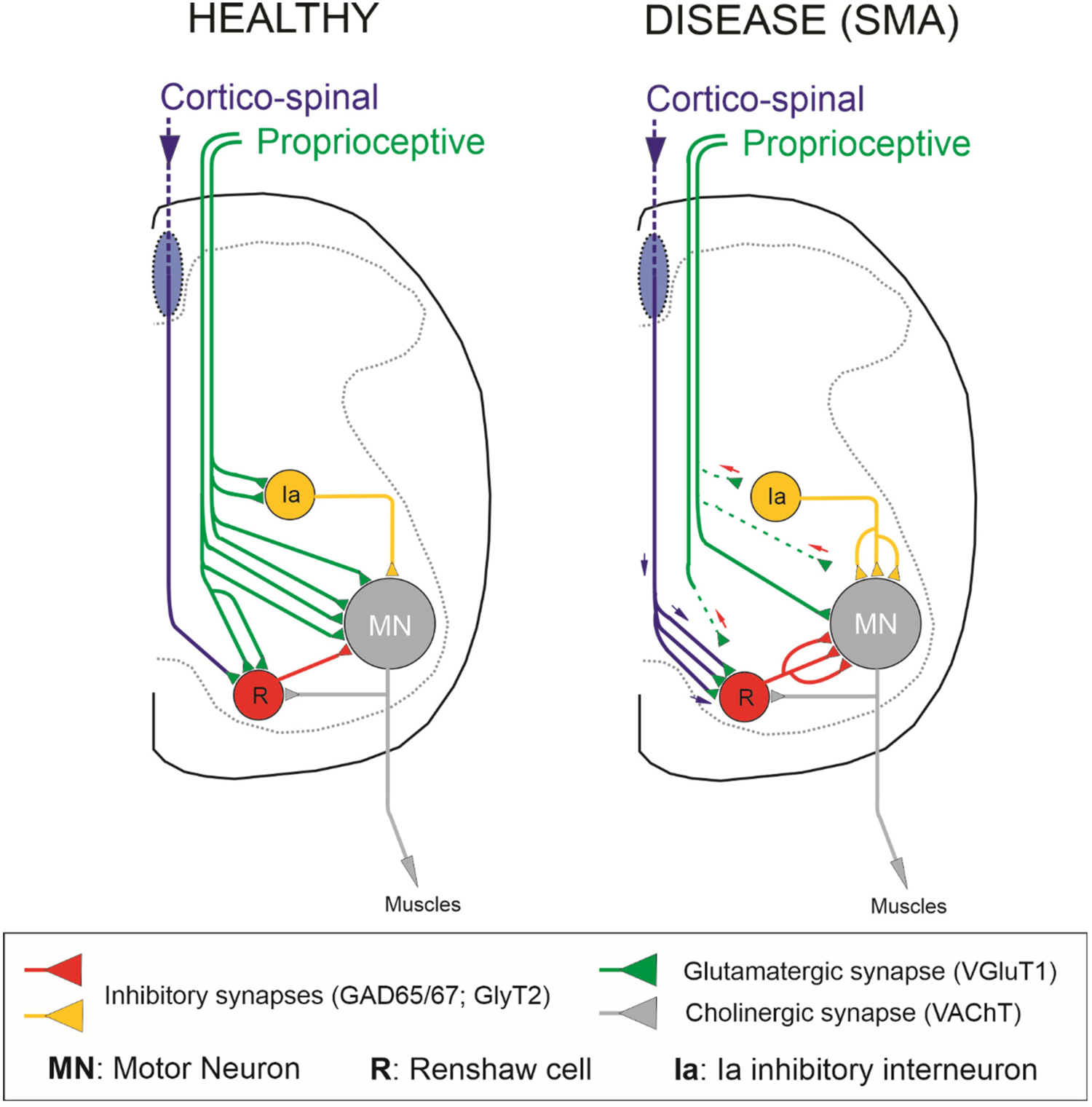
Neuronal circuit changes conferring an increase in tonic inhibitory drive on motor neurons in a severe SMA mouse model. Under healthy conditions, lumbar motor neurons (grey) receive excitatory-glutamatergic synaptic drive from proprioceptive fibers (green) and inhibitory synapses from Renshaw cells (red) and Ia inhibitory interneurons (yellow). In SMA, vulnerable motor neurons receive decreased proprioceptive synaptic drive while their inhibitory synapses from Renshaws and Ia inhibitory interneurons increase. Furthermore, Renshaw cells receive higher than normal excitation from corticospinal glutamatergic synapses (blue/green).

### Aberrant increase in inhibition is a maladaptive mechanism for motor deficits in SMA

For normal function to occur, individual neurons and neuronal circuits must maintain a balanced E/I while synaptic activity is fluctuating or perturbed (He *et al*., 2016; Zhou *et al*., 2014). At the single neuron level, it has been reported that reduced excitatory synaptic transmission decreases inhibitory synaptic puncta and mIPSCs (He *et al*., 2018). However, reductions in inhibitory synaptic drive do not lead to a corresponding decrease in excitatory inputs (Shen *et al*., 2011). Thus, it is thought that E/I balance is not a bidirectional neuronal response to any disturbance of either excitatory or inhibitory synapses, but it is rather dominated by changes in excitation (He and Cline, 2019). Intriguingly, our study shows that reduced excitatory synaptic drive of SMA mice motor neurons do not result in homeostatic compensations at inhibitory synapses. On the contrary, inhibitory synapse numbers and strengths on motor neurons are paradoxically increased. Demonstration that this response is maladaptive comes from “rescue” experiments in which knockdown of gephyrin improves the cellular, circuit and behavioural phenotype of SMA mice. Inhibitory synapse strengths follow the maturation of local excitatory inputs strengths in spinal cord sensory-motor circuits (Allain et al., 2011; Delpy et al., 2008; González-Forero and Alvarez, 2005) and this normal process might be altered by SMA pathology. One candidate mechanism is excessive activity in inhibitory interneurons due to their higher input resistance, lower action potential thresholds and possible alternate excitatory inputs. In other brain regions, activity-dependent transcription factors like Npas4 regulate the number and strength of synapses formed by inhibitory interneurons (Lin et al., 2008; Spiegel et al., 2014) raising the possibility that increased firing in spinal inhibitory interneurons in SMA could drive inhibitory synaptogenesis and augment synaptic strengths. Alternatively, during embryonic periods, in which GABA and glycine are depolarizing (Delpy *et al*., 2008), homeostatic synaptic plasticity on motor neurons is driven by excitatory GABAergic synapses (Wenner, 2014) and early deficiencies in GABA/glycine interneurons may cause abnormal synaptic organizations and strengths on target motor neurons in SMA.

### Deficits in motor neuron output due to unwarranted dysregulation of recurrent and reciprocal inhibitory spinal circuits in SMA

Motor output and muscle activity depend on motor units’ recruitment and firing rates. Recruitment gain is the relationship between the intensity of the synaptic drive to a motor pool and its output (Hultborn *et al*., 2004; Kernell and Hultborn, 1990). Experiments in the cat have demonstrated that variations in the distribution of synaptic inputs to different types of motor units can change their recruitment gain (Nielsen et al., 2019). One prominent inhibitory interneuron proposed to control output across spinal motor neuron pools is the Renshaw cell (Hultborn et al., 1979), which is responsible for recurrent inhibition (Eccles *et al*., 1954; Renshaw, 1946). The input/output relation across a motor neuron pool is determined by the intrinsic properties of motor neurons as well as their synaptic input, and there is evidence that recurrent inhibition is effective in reducing synaptically-induced motor neuron firing rates (Hultborn *et al*., 2004). An additional source of inhibition in motor neurons is from Ia inhibitory interneurons, which are responsible for reciprocal inhibition of antagonistic motor neurons (Eccles et al., 1956; Hultborn, 1972; Jankowska, 1992). Recurrent and reciprocal inhibition are functional in neonatal mice (Bhumbra *et al*., 2014; Moore *et al*., 2015; Sapir et al., 2004; Wang et al., 2008; Zhang *et al*., 2014) and human newborns (Mc Donough et al., 2001). Importantly, both recurrent and reciprocal inhibitory effective synaptic currents distribute uniformly within a pool of motor neurons (Binder et al., 2002) and their synapses are located on cell bodies and proximal dendrites at short electronic distances (Burke et al., 1971; Fyffe, 1991; Worthy *et al*., 2023). Thus, they exert effective modulation of motor neuron firing, even in the early postnatal spinal cord, by powerfully shunting the proximal somato-dendritic membrane in regions spatially close to action potential trigger zones (Bhumbra *et al*., 2014). Excessive synaptic activity at the level of cell body and originating in both inhibitory interneuron types could therefore effectively reduce recruitment and maximal firing rates of motor neurons under SMN deficiency. Vulnerable SMA motor neurons receiving reduced excitation (Fletcher *et al*., 2017; Simon *et al*., 2016) will be substantially more difficult to be recruited under the undue influence of increased inhibitory drive from these key premotor inhibitory interneurons.

### Opposing neuronal circuit mechanisms act in SMA and ALS: therapeutic implications

Insights into disease mechanisms can be drawn by comparing two most prominent motor neuron diseases, SMA and ALS. Studies in ALS, reveal that motor units increase their excitability, evident by an increase of fasciculation potentials, double discharges of motor units (Kostera-Pruszczyk et al., 2002; Piotrkiewicz et al., 2008), and aberrant single motor unit firing (Piotrkiewicz *et al*., 2008). Furthermore, glutamatergic neurotransmission through sensory afferents is affected and negatively impacts motor neuron function in mouse models of ALS (Bączyk et al., 2020; Seki et al., 2019), while decreased expression of VGluT2 (a vesicular transporter for glutamatergic neurotransmission) in SOD1^G93A^ mice reduced motor neuron loss but had no impact on disease onset or life span (Wootz et al., 2010). Reduction of excitatory synapses from Ia proprioceptive fibers in SOD1^G93A^/Egr3^-/-^ mice also slowed down motor neuron loss, but again did not alter disease progression (Lalancette-Hebert et al., 2016). This is in contrast to the SMA, in which deficits in Ia proprioceptive synapses is an early and major contributor to disease phenotype (Fletcher *et al*., 2017; Mentis *et al*., 2011). Recurrent inhibition is abnormally reduced in ALS patients (Özyurt et al., 2020; Raynor and Shefner, 1994) and synapses forming the recurrent inhibitory circuit degenerate around the time motor symptoms start in the SOD1^G93A^ mouse model (Wootz et al., 2013). In striking contrast, our current study in SMA demonstrates that recurrent inhibition is aberrantly increased on SMA motor neurons. In ALS, reciprocal inhibition measured through the Hoffman H-reflex is also impaired in patients (Misra and Kalita, 1998) as well as rate-dependent depression of H-reflexes indicating overall disinhibition of motor circuits modulating motor unit responses to the monosynaptic Ia reflex (Zhou et al., 2022). Disinhibition of motor neurons in ALS has been reported with a loss of perisomatic synapses from V1 interneurons and the V1 interneurons themselves, including Foxp2+ interneurons (Allodi *et al*., 2021; Salamatina *et al*., 2020). Additionally, previous studies reported a decrease in glycine receptor expression in postmortem tissues of ALS patients (Hayashi et al., 1981; Whitehouse et al., 1983), observations that were recapitulated in mouse models as evidenced by a reduction of GlyT2 and GAD65/67 expression in ventral horns of SOD1^G93A^ mice (Hossaini et al., 2011). In contrast, our study shows the Foxp2 interneurons which are involved in reciprocal inhibition show similar dysfunctional changes to those observed in Renshaw cells in SMA mice. Thus, while dysregulation of inhibitory inputs to motor neurons is pathogenic highlighting the central role of inhibitory circuit dysfunction in motor disease, opposite changes in the inhibitory control of motor neurons in ALS compared to SMA may help explain the different motor symptoms in the two diseases. In ALS, we observe hyperreflexia, excessive co-contractions, cramps and muscle fasciculations, while in SMA, patients exhibit hyporeflexia and overall muscle paralysis with diminished motor output (SMA newborns). Importantly, we provide proof-of-concept that either genetic or pharmacological approaches targeting excess inhibitory drive through SMN independent mechanisms can be therapeutically relevant in SMA.

### Abnormal increase in excitation of spinal Renshaw cells by cortico-spinal synapses in SMA

What drives the increase in inhibitory drive on SMA motor neurons? In adult animals, skilled movement depends on the coordination of control signals from descending pathways and afferent fibers. Two of the most critical signals on motor circuits are the descending cortico-spinal and the peripheral sensory proprioceptive afferents. Both signals mature during early postnatal development with the proprioceptive slightly ahead of the cortico-spinal circuitry (Martin et al., 2007). We have previously demonstrated that proprioceptive fibers contact Renshaw cells monosynaptically in neonatal mice (Mentis *et al*., 2006), an effect that occurs also in embryonic chick spinal cords (Wenner and O’Donovan, 1999). In most animals, cortico-spinal synapses do not contact lumbar motor neurons directly, whereas both cortico-spinal and proprioceptive axons synaptically converge onto common spinal interneurons, including inhibitory ones (Chakrabarty et al., 2009; Hultborn and Santini, 1972; Jankowska and Edgley, 2010). Intriguingly, cortico-spinal and proprioceptive afferent synapses compete such that the loss of one input induces the expansion of the other (Chakrabarty and Martin, 2011; Jiang *et al*., 2016; Tan et al., 2012). To this end, we show that dysfunctional proprioceptive synapses on Renshaw cells are progressively eliminated, and their place taken over by cortico-spinal synapses. It is, therefore, logical to speculate that sensory synapses dysfunction will likely result in abnormal excitation of Renshaw cells and Ia inhibitory interneurons by alternate inputs. Together with increased inhibitory interneuron excitability and the possibility that these interneurons are spontaneously active, they could impose a disproportionate inhibitory synaptic drive on vulnerable SMA motor neurons, rendering them more difficult to be recruited (Fig. 8). Thus, we propose that synaptic competition between cortico-spinal and proprioceptive synapses on Renshaw and Ia inhibitory pre-motor interneurons is a key synaptic mechanism that contributes to neuronal circuit dysfunction and the progressive loss of motor control, resulting in reduced motor neuron output and eventual muscle paralysis in SMA.

## METHODS

### Animals and genotyping

All surgical procedures were performed on postnatal mice in accordance with the National Institutes of Health (NIH) Guidelines on the Care and Use of Animals and approved by the Columbia animal care and use committee (IACUC). Animals of both sexes were used in this study. The original breeding pairs for the SMA mice used in our study (Smn^+/−^/SMN2^+/+^/SMNΔ7^+/+^) were purchased from Jackson Mice (Jax stock #005025; FVB background). Tail DNA PCR genotyping protocols for SMA-Δ7 mice were followed as described on the Jackson website (www.jax.org).

To restore SMN selectively in proprioceptive neurons, we used a mouse model of SMA harboring a single targeted mutation and two transgenic alleles, resulting in the genotype Smn^Res/+^;SMN2^+/+^;SMNΔ7^+/+^ (where *Smn* is used for the mouse *Smn1* gene and *SMN* for the human *SMN2* gene). The allele carrying the targeted mutation (Smn^Res^) is engineered to revert to a fully functional Smn allele upon Cre-mediated recombination (Cre^+/−^;Smn^Res/−^;SMN2^+/+^;SMNΔ7^+/+^). *SMN2* is the human gene and SMNΔ7 corresponds to the human SMN cDNA lacking exon 7. In the absence of the Cre recombinase (Cre^−/−^;Smn^Res/−^;SMN2^+/+^;SMNΔ7^+/+^) the phenotype of these mice is similar to that of the SMNΔ7 SMA mice. Restoration of SMN protein in proprioceptive neurons was achieved by crossing the conditional inversion SMA mice with Pv^Cre^ mice (Jax stock #008069), which express Cre under the control of the parvalbumin (Pv) promoter. Parvalbumin is expressed exclusively in proprioceptive neurons during the first 10 postnatal days and was expressed similarly in WT and SMA mice (Fletcher *et al*., 2017).

### Behavioural analysis

Mice from all experimental groups were monitored daily, weighed, and three righting reflex tests were timed and averaged as described previously (Fletcher *et al*., 2017). Mice with 25% weight loss and an inability to right were euthanized with carbon dioxide to comply with IACUC guidelines. Righting time was defined as the time for the pup to turn over after being placed completely on its back. The cut-off test time for the righting reflex was 60 secs to comply with IACUC guidelines.

### Transection experiments

Experiments were conducted on both wild type and SMA mice at P8. Pups were anesthetized with isoflurane (5% induction and 2.5% maintenance). A transverse fine slit was opened through the vertebral column with a pair of forceps at the T4/5 spinal segment. The spinal cord transection was made with a pair of fine scissors. The skin was sutured and the pups were returned to their cage following recovery from anesthesia. The success of the transection was validated by the lack of responses to hindlimb muscles after a light tail pinch. At P10, animals were deeply anesthetized and transcardially perfused with 4% paraformaldehyde (PFA) and the spinal cord was removed. Following an overnight fixation, the spinal cord was embedded in 5% agar and sectioned into 75 μm transverse sections using a vibratome. In addition, the extent of the transection and level of the transection were verified. Only animals with complete transection at the L4/5 spinal segments were included in the study.

### Physiology using the intact neonatal *ex vivo* spinal cord preparation

#### Current clamp recordings

Experimental protocols used in this study have been described before (Fletcher *et al*., 2017; Mentis *et al*., 2011). Animals were decapitated and the spinal cords dissected and removed under cold (∼12°C) artificial cerebrospinal fluid (aCSF) containing in mM: 128.35 NaCl, 4 KCl, 0.58 NaH_2_PO_4_.H_2_0, 21 NaHCO_3_, 30 D-Glucose, 1.5 CaCl_2_.H_2_0, and 1 MgSO4.7H_2_0. The spinal cord was then transferred to a customized recording chamber placed under the objective of an epifluorescent (Leica DM6000FS) microscope. The preparation was perfused continuously with oxygenated (95% O_2_ / 5% CO_2_) aCSF (∼10 ml/min). Ventral roots and dorsal roots were placed into suction electrodes for stimulation or recording.

Whole-cell recordings were performed at room temperature (∼21°C) and obtained with patch electrodes advanced through the lateral or ventral aspect of the spinal cord. Patch electrodes were pulled from thin-walled borosilicate glass capillary with filament (Sutter Instruments) using a P-1000 puller (Sutter Instruments) to resistances between 5–8 MΩ. The electrodes were filled with intracellular solution containing (in mM): 10 NaCl, 130 K-Gluconate, 10 HEPES, 11 EGTA, 1 MgCl_2_, 0.1 CaCl_2_ and 1 Na_2_ATP, 0.1 Cascade Blue hydrazide (Life Technologies), and in some experiments with 0.5 mg/ml Neurobiotin (Vector Labs). pH was adjusted to 7.2–7.3 with KOH (the final osmolarity of the intracellular solution was 295–305 mOsm). Motor neurons, Renshaw cells, putative Ia inhibitory interneurons and other unidentified spinal neurons were targeted blindly. The identity of recorded neurons as motor neurons was confirmed by evoking an antidromic action potential by stimulation of the cut ventral root. Renshaw cells were identified physiologically by the occurrence of evoked graded excitatory synaptic potentials after ventral root stimulation. Ia inhibitory interneurons were putatively characterized as such because of their relative position (dorsal to the motor neuron nucleus), monosynaptic activation by proprioceptive fibers following dorsal root stimulation, but no appreciable response following ventral root stimulation. Other unidentified spinal interneurons were group together as neurons that did not respond monosynaptically from either dorsal root or ventral root stimulation. All neurons were accepted for further analysis only if the following three criteria were met: (i) stable resting membrane potential of −50 mV or more negative (ii) an overshooting action potential following current injection and (iii) at least 30 mins of recording.

For the measurements of passive membrane properties, neurons were injected with sequential steps of negative and positive currents for 100 ms in small steps of current at −60 mV membrane potential. The input resistance (MΩ) was calculated from the slope of the current/voltage plot within the linear range. Membrane time constants (ms) were calculated as 63% of the maximal negative amplitude during the application of the current pulse. The membrane capacitance (MΩ/ms) of each cell was calculated by dividing the input resistance by the time constant. Measurements were taken from an average of 3 sweeps. Spontaneous activity from wild type and SMA motor neurons and Renshaw cells was measured at their own resting membrane potential (RMP). The frequency of spontaneous activity (Hz) was calculated from a 1-minute recording. To compare statistically the firing frequency in all experimental groups we used small steps of current (10 pA) above the minimum current required to elicit repetitive firing for 1 sec. The firing frequency (Hz) was calculated using the event detection function in Clampfit.

Synaptic potentials were recorded from individual Renshaw cells (DC - 3 kHz, Multiclamp 700B, Molecular Devices) in response to a brief (0.2 ms) orthodromic or antidromic stimulation (A365, current stimulus isolator, WPI, Sarasota, FL) of the dorsal root or ventral root respectively (L2 or L3). The stimulus threshold was defined as the current at which the minimal evoked response was recorded in 3 out of 5 trials. The dorsal or ventral root was stimulated at different multiples of threshold. Recordings were fed to an A/D interface (Digidata 1440A, Molecular Devices) and acquired with Clampex (v10.2, Molecular Devices) at a sampling rate of 10 kHz. Data were analyzed off-line using Clampfit (v10.2, Molecular Devices). The monosynaptic component of the EPSP amplitude was measured from the onset of response to 3 ms. Measurements were taken from averaged traces of 5 trials elicited at 0.1 Hz. Bridge balance was applied to all recordings. The liquid junction potential was calculated as −5 mV but was not corrected. Measurements were made on averaged traces (3 – 5 trials).

γ (gamma) motor neurons were not included in our analysis. γ motor neurons were identified by the presence of an antidromic action potential, but lack of direct monosynaptic activation from proprioceptive sensory fibers.

#### Voltage clamp recordings

Whole-cell voltage-clamp recordings were obtained from antidromically identified motor neurons in the L1 and L2 spinal segments. Motor neurons were targeted “blindly” either from the lateral or ventral aspect of the spinal cord. Patch electrodes contained the following (in mM): 120 CsCl, 4 NaCl, 4 MgCl_2_, 1 Cl_2_Ca, 10 HEPES, 0.2 EGTA, 3 Mg-ATP, and 0.3 GTP-Tris. In some of the experiments, 1% Neurobiotin (Vector Laboratories) was added to the internal solution. Only recordings with access resistance less than 20 MΩ were included in our analysis. The access resistance was checked throughout the experiments and recordings were abandoned if it changed more than 15%. Neurons were voltage clamped at −75 mV. Synaptic currents were recorded and low-pass bessel filtered at 5 kHz with an Multiclamp 700B amplifier. Data were digitized at 10 kHz and acquired using Clampex (v10.2, Molecular Devices). For each motor neuron, we obtained approximately 2 mins of continuous recording of spontaneous activity under drug combinations that pharmacologically isolated the synaptic currents of interest.

To isolate miniature spontaneous synaptic currents of GABAergic and/or glycinergic origin [miniature inhibitory postsynaptic currents (mIPSCs)], recordings were performed in the presence of tetrodotoxin (TTX, 1 μM; Alomone Labs), the glutamate receptor blockers 6-cyano-7-nitroquinoxaline-2,3-dione (CNQX, 10 μM; Sigma) and 2-amino-5-phosphonovaleric acid (APV, 100 μM, Sigma), as well as the cholinergic receptor blockers, mecamylamine (50 μM, Sigma), dihydro-β-erythroidine (dHβE, 50 μM, Sigma) and D-tubocurarine chloride (30 μM; Sigma), similar to a previous report (González-Forero and Alvarez, 2005). All drugs were applied to the bath solution. Glycinergic and GABAergic miniIPSCs were verified physiologically since they were abolished following the addition of bicuculline methiodide (10 μM; Sigma) and strychnine hydrochloride (0.25 μM; Sigma) to the bath solution.

#### *In vivo* EMG recordings

Electromyography (EMG) was performed in P11 mice. The EMG electrode was placed on the Quadratus Lumborum under anaesthesia, induced by 5% isoflurane and maintained by 1.5-2% during the electrode implantation. The electrode was bipolar and made of two silver Teflon-coated wires. The iliopsoas muscle was identified following a small incision from the left side in the stomach area, making sure that the peritoneum was not punctured. The naked tips of the bipolar electrode were bent to ensure that the electrode will remain in place following their insertion into the muscle. Correct placement of the electrode into the iliopsoas was verified by muscle contraction following a brief (0.2 ms) stimulation with an isolated current stimulator (A365, current stimulus isolator, WPI). The pup was allowed to recover from anaesthesia for approximately 30mins. At this time point, the pup was placed on a warm surface area, the EMG electrode was connected to a pre-amplifier (10x amplification) and the signal was further amplified to a final 1K amplification (Digidata 1440A, Molecular Devices). Recordings were acquired with Clampex (v10.2, Molecular Devices) at a sampling rate of 10 kHz. Data were analyzed offline using Clampfit (v10.2, Molecular Devices). The pup was placed on its back and allowed to right itself while EMG recordings were acquired. This was repeated at least three times to ensure consistency and acquisition of high-quality recordings. At the end of the EMG recordings, the pup terminally anaesthetized, transcardially perfused with 4% paraformaldehyde and the spinal cord was removed for further examination using immunohistochemistry.

#### Immunohistochemistry

Detailed protocols for immunohistochemistry used in this study have been previously described (Fletcher *et al*., 2017; González-Forero and Alvarez, 2005; Mentis *et al*., 2011). Antibodies used in this study are listed in Table 1. Mouse spinal cords were either i) transcardially perfused with 4% paraformaldehyde followed by overnight post fixation in 4% paraformaldehyde or, ii) immersion fixed overnight in 4% formaldehyde diluted in PBS. L1 – L2 segments were either embedded in warm 5% Agar for cutting serial transverse sections on a Vibratome (75 μm thickness) or cut frozen on Peltier stage of a sliding freezing microtome (50 μm thick sections) or in cryostat (25 μm thick). Sections were blocked with 10% normal donkey serum in 0.01M PBS with 0.1% Triton X-100 (PBS-T; pH 7.4) and incubated overnight at room temperature in different combinations of antisera in PBS-T. For experiments involving anti-mouse antibodies, sections were pre-incubated for 1 hour in M.O.M blocker (Vector Laboratories) in PBS-T to block endogenous antigens. The following day, sections were washed in PBS-T and secondary antibody incubations were performed for 3 hours with the appropriate species-specific antiserum diluted in PBS-T. Sections were subsequently washed in PBS, mounted on glass slides using Vectashield (Vector Laboratories). The secondary antibodies (Jackson Labs) used in this study were: Alexa 488, Cy3, and Cy5 (dilution 1:250).

**Table 1:**
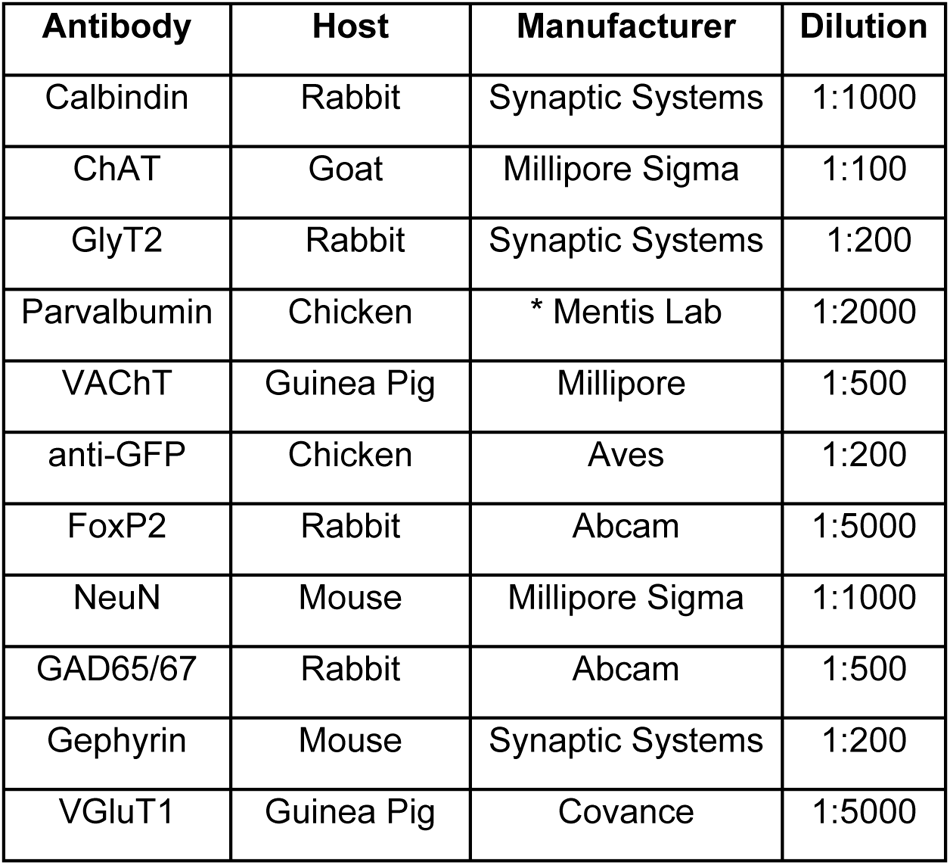
Primary antibodies used in this study.

In some experiments, a quadruple type of staining (four fluorochromes) was performed using the following protocol. In these experiments section preparation and incubation in primary antibodies was as above but selected immunoreactivities were revealed using a three-step procedure. First the primary antibody was detected with donkey anti-goat biotinylated antibody (according to the chosen primary antibody) following the incubation with primary antibodies in blocking serum for 3 hours. Subsequently, the sections were washed with PBS-Tr six times for 10 min each time and then we added Streptavidin–Alexa 405 (Jackson Labs) to detect the immunoreactivities in the “blue” channel.

#### Confocal microscopy and image analysis

Quadruple immunofluorescence (405, 488, 555 and 657 excitation wavelengths) were visualized with a Fluoview FV1000 laser-scanning confocal microscope (Olympus, Japan) or with Leica SP5 or Leica SP8 confocal microscopes (Leica, Germany). Sections were analysed using Leica or Olympus or Neurolucida software (MBF Biosciences). For all immunohistochemical analysis, at least three animals from each postnatal stage were used (with one exception: comparison of VGluT1 and VAChT synaptic densities of P4 Renshaw cells, see below). Analysis was performed from single optical plane images acquired with an ×63 oil objective at 4096 × 4096 dpi resolution using an SP5 Leica confocal microscope. Only motor neuron somata (identified by ChAT immunoreactivity) in which the nucleus was present were included in the analysis. Gephyrin or GlyT2 or GAD65/67 density was calculated by motor neuron soma circumference, divided by the number of positive gephyrin clusters or GABAergic or glycinergic synapses.

#### Synaptic density of proprioceptive and cholinergic inputs onto Renshaw cells

Lumbar (L1-L2) spinal cord segments from postnatal (P3-P11) mice were sectioned in the transverse plane, 25µm thick, on a freezing cryostat and collected directly on slides for immunostaining with different combination of calbindin, VGluT1, parvalbumin and VAChT antibodies using the procedures described before (Table 1). Immunoreactive sites were revealed with species-specific secondary antibodies coupled to different fluorochromes (Alexa 488 or FITC, CY3, Alexa 647 or DyLight647 at 1:100).

Calbindin positive cells found in the ventral horn of the spinal cord close to the exit of motor neurons axons from the grey matter and close to the ventral root, which received synapses from motor neuron axon collateral were defined as Renshaw cells. These cells were imaged on an Olympus FV1000 confocal microscope or Leica SP5 or Leica SP8. To reconstruct VGluT1 and VAChT coverage on Renshaw cells, optical section stacks (step size, 0.5 μm) were captured throughout the cell body and proximal dendrites of the neurons using a 60x oil immersion objective (numerical aperture, 1.4) and digitally zoomed (x1.5). Confocal image stacks were uploaded into Neurolucida (MBF Bioscience), where soma and proximal dendritic arbors were reconstructed in 3D. Synaptic contacts that were either VGLUT1/Parvalbumin^+^ or VAChT^+^ were counted and marked along cell body and the dendritic arbor for analysis. The somatic, linear, and surface area densities were then calculated based on these reconstructions and compared between WT and SMA. Images used for figure composition were filtered (high-Gauss filter, Image Pro-Plus 4.0; Media Cybernetics) and adjusted for contrast, brightness, and dynamic resolution for best quality presentation without changing or altering the information content in the images.

#### Labeling of motor neuron axon collaterals

In some experiments, we labelled motor neuron axon collaterals by retrograde tracing of a fluorescent tracer using the *ex vivo* spinal cord preparation. Following dissection and removal of the spinal cord from the vertebral column from neonatal mice, the L1 ventral root was placed inside a suction electrode and backfilled with a fluorescent dextran to fill the somato-dendritic tree of motor neurons including the axon collaterals. The spinal cord was perfused with cold (∼10 °C), oxygenated (95% O_2_, 5% CO_2_) aCSF (containing, in mM, 128.35 NaCl, 4 KCl, 0.58 NaH_2_PO_4_, 21 NaHCO_3_, 30 d-glucose, 0.1 CaCl_2_ and 2 MgSO_4_). After 12–16 h the spinal cord was immersion-fixed in 4% paraformaldehyde and washed in 0.01 M PBS. Sections were subsequently processed for immunohistochemistry.

#### *In vivo* retrograde labeling of muscle-identified (iliopsoas) motor neurons

Motor neurons supplying the *iliopsoas* (IL) and *quadratus lumborum* (QL) muscles were retrogradely labeled *in vivo* by intramuscular injection of CTb conjugated to Alexa 488. Newborn (P0) mice were anesthetized by isoflurane inhalation. A small incision in the left iliac (inguinal) area was made to access the IL/QL muscles, taking care not to puncture the peritoneum. The muscles were injected with ∼1 μl of 1% CTb-Alexa 488 in PBS using a finely pulled glass micropipette. The CTb was delivered by pressure to an adapted micro-syringe. The incision was closed with sutures. The spinal cord was taken at P11 following verification by fluorescence of accurate injection of CTb in the muscles and processed for immunohistochemistry.

#### AAV9 vectors

For *in vivo* knockdown of gephyrin by RNAi a self-complementary vector containing AAV2 ITRs was engineered by standard molecular biology methods to harbor a mouse U6 promoter driving expression of an shRNA targeting the sequence “CCCTTCTTAGTATGCTTCA” of mouse Gephyrin as well as a CMV promoter driving GFP expression (AAV9-Gephyrin_RNAi_). Production and purification of AAV9 vectors was carried out as previously described (Simon *et al*., 2017). Titering was done using cybergreen qPCR using previously described primers against GFP (Simon *et al*., 2017). An additional titering method using Quantiflour ds DNA system (Promega) was performed according to manufacturer’s instructions. AAV9-Gephyrin_RNAi_ was administered by intracerebroventricular (I.C.V.) injection. Wild type (controls) and SMA mice were injected at P0. The dosage of injection was 5.2*10^10 genome copies per animal. As a control for AAV9-Gephyrin_RNAi_, mice were injected with AAV9-GFP (Simon *et al*., 2019). Righting reflex time and body weight were monitored daily. The lifespan of treated mice was also monitored and recorded.

#### *In vivo* daily treatment with Org-25543

Wild type and SMA mice were used for this experiment. For systemic administration of Org-25543 (Tocris), P0 – P7 pups were injected daily subcutaneously with a dose of 1.5 mg/kg dissolved in saline and monitored for body weight and righting times until P7. The righting was performed prior to the injection of the drug, three times and averaged.

#### Statistics

Results are expressed as means ± s.e.m. Statistical analysis was performed using GraphPad Prism 6. Comparison was performed by either Student’s t-test or one-way ANOVA (post hoc comparison methods are indicated in the figure legends when necessary). Results were considered statistically significant if P < 0.05. The D’Agostino and Pearson omnibus normality test was used to assess the normality for all data. If violated, non-parametric tests were used. No statistical methods were used to predetermine sample sizes, but our sample sizes are similar to those reported in previous publications. No randomization was used. Data collection and analysis were not performed blind to the conditions of the experiments. Statistical comparison was performed on the average value from individual mice. In physiological experiments, a single Renshaw cell, or Ia inhibitory interneuron, or an unidentified interneuron was recorded from a single mouse.

## ACKNOWLEDGMENTS

We thank Dr Marco Capogrosso for providing critical comments to our study. This work is supported by R01-NS078375 (GZM), R01-NS125362 (GZM), R01-AA027079 (GZM), The SMA Foundation (GZM), Project ALS (GZM); R01-NS102451 (LP), R01-NS114218 (LP), R01-NS116400 (LP); R01 NS 047357 (FJA).

## AUTHOR CONTRIBUTIONS

EVF, FJA and GZM conceived the study; EVF, JIC, TMR, JGP, MVA, NS performed experiments; EVF, FJA, JIC, TMR, LP, GZM analyzed the data; MVA, LP designed viruses, JER produced viruses, EVF, FJA and GZM wrote the paper with contributions from all authors.

## SUPPLEMENTAL FIGURES

**Suppl. Fig. 1.**
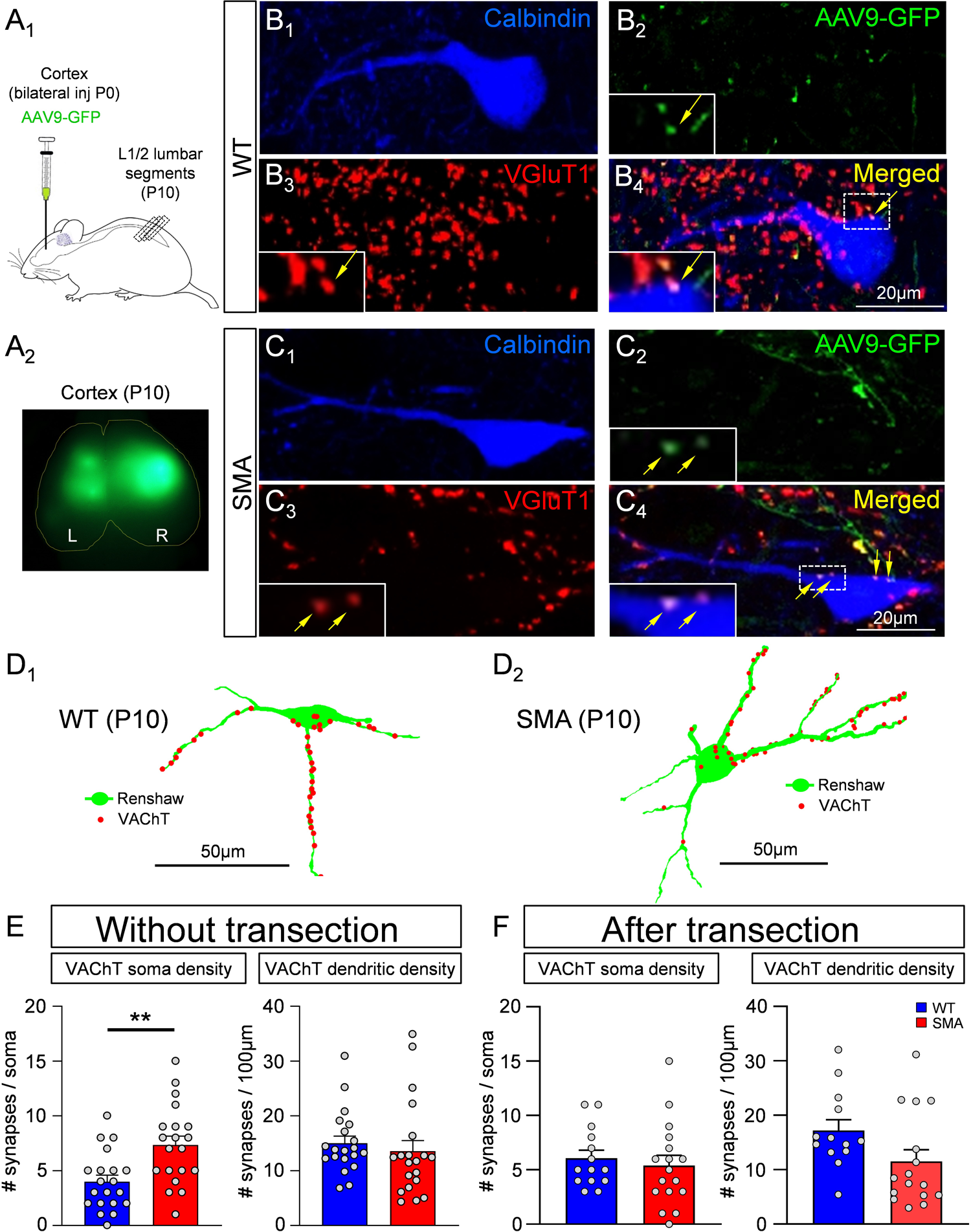
(Associated with Fig. 3). Renshaw cells receive VGluT1+ synapses originating in corticospinal neurons. (A_1-2_) Experimental protocol for labelling corticospinal synapses (A_1_). Mice injected at birth (P0) with AAV9-GFP bilaterally in the cortex (A_2_). At P10, the L1 and L2 spinal segments were examined with immunohistochemistry. **(B_1-4_, C_1-4_)** Single plane confocal images of a wild type (B_1-4_) and a SMA (C_1-4_) Renshaw cell labelled with calbindin (blue, B_1_ and C_1_), AAV9-GFP (green, B_2_ and C_2_), VGluT1 (red, B_3_ and C_3_) antibodies. Merged images are shown in B_4_ and C_4_. Insets are areas indicated by the dotted boxes, showing GFP+ and VGluT1+ synapses on the soma (yellow arrows) of Renshaw cells. **(D_1,2_)** Neurolucida reconstruction of a wild type (D_1_) and a SMA (D_2_) Renshaw cell with cholinergic (VAChT+) synapses marked by red dots. **(E)** Number of cholinergic (VAChT+) synapses on the soma (left graph) and dendrites (right graph) of Renshaw cells in wild type (blue) and SMA (red) without spinal cord transection at P10. Differences were significant on cell bodies (** p=0.0018 unpaired two-tailed t-test) but not on dendrites. (n=20 or 10 Renshaw cells per animal; N=2 WT and 2 SMA mice) **(F)** Number of cholinergic (VAChT+) synapses on the soma (left graph) and dendrites (right graph) of Renshaw cells in wild type (blue) (n=14, N=3) and SMA (red) two days after T4 spinal cord transection at P10 (n=17, N=3). Differences are non-significant (unpaired two-tailed t-test).

**Suppl. Fig. 2.**
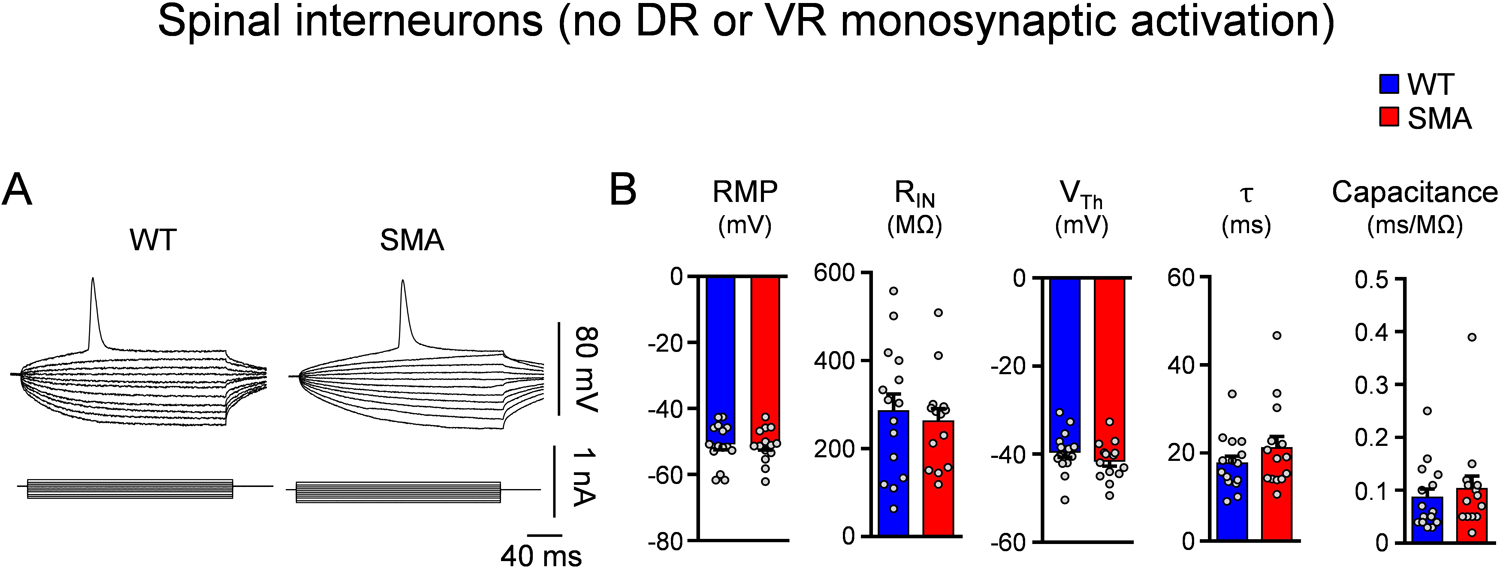
(Associated with Fig. 4). No electrophysiological differences in SMA spinal interneurons that do not receive proprioceptive synapses. (A) Superimposed voltage responses (top traces) following current injection (bottom traces) in spinal interneurons that do not receive direct proprioceptive synapses in wild type and SMA mice at P4. (B) Resting membrane potential (RMP), input resistance (R_IN_), voltage threshold (V_Th_), time constant (τ) and capacitance of spinal interneurons without direct proprioceptive activation in wild type (blue, n=15 neurons, N=15 mice) and SMA (red, n=14 neurons, N=14 mice) mice at P4.

**Suppl. Fig. 3.**
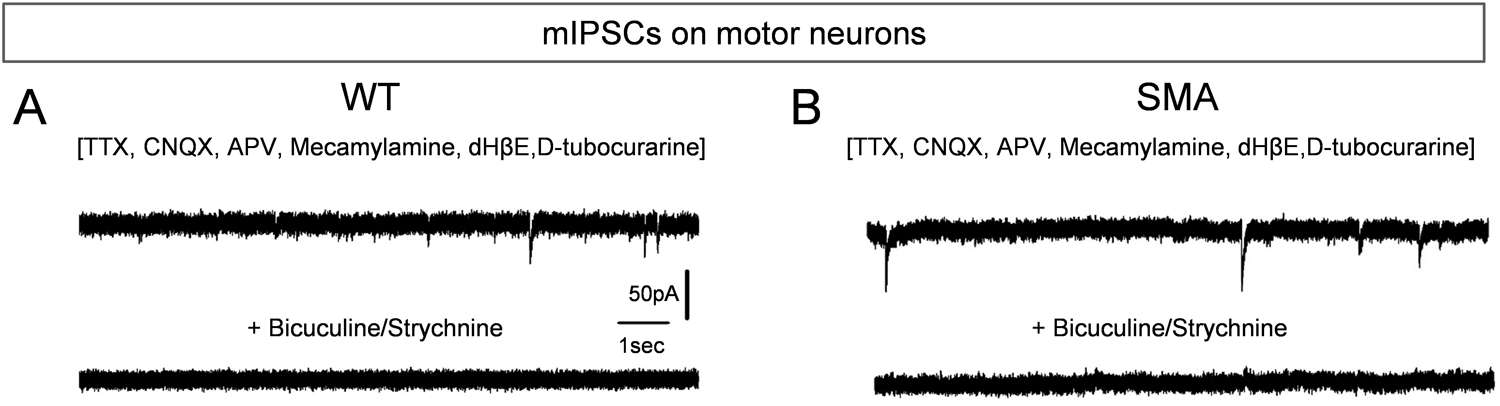
(Associated with Fig. 5). Validation of mIPSCs in wild type and SMA motor neurons. Current recordings from voltage clamp experiment in wild type **(A)** and SMA **(B)** motor neurons in which mIPSCs (top traces) were abolished by application of bicuculine and strychnine (bottom traces).

**Suppl. Fig. 4.**
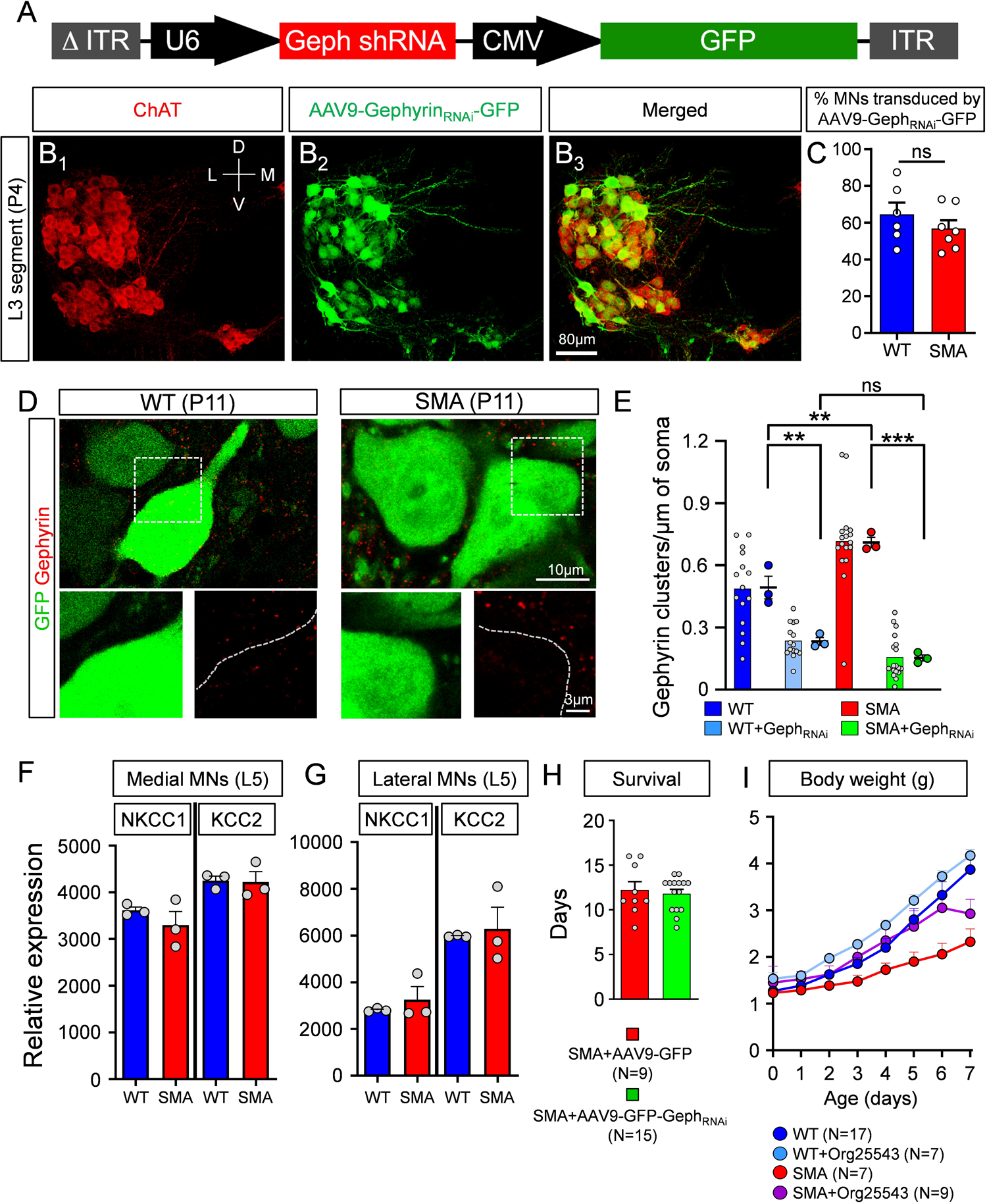
(Associated with Fig. 6). Validation of gephyrin knockdown; no difference in NKCC1 or KCC2 between wild type and SMA mice; behavioral phenotype injected with AAV9-Geph_RNAi_ or treated *in vivo* with Org25543. (A) Map of the plasmid for gephyrin knockdown. **(B_1-3_)** Confocal images from the ventral spinal cord of a wild type mouse at P4 showing ChAT (red, B_1_), AAV9-Gephyrin_RNAi_-GFP (green, B_2_) immunoreactivity and their merged image (B_3_). **(C)** Percentage of motor neurons (MNs) transduced by AAV9-Gephyrin_RNAi_-GFP in wild type (n=295 MNs, N=6) and SMA (n=191 MNs, N=7) mice at P11. Each data point represents one mouse. **(D)** GFP (green) and gephyrin (red) in a wild type (left) and a SMA (right) motor neuron at P11. Images at the bottom are higher magnification areas from the dashed boxes, respectively. **(E)** Number of gephyrin clusters per μm of motor neuron membrane in wild type mice (blue; n=15 MNs, N=3 mice), wild type mice injected with AAV9-Geph_RNAi_ (cyan; n=15 MNs, N=3 mice), SMA mice (red; n=17 MNs, N=3 mice) and SMA mice injected with AAV9-Geph_RNAi_ (green, n=18 MNs, N=3 mice). ** p=0.002, WT vs WT+Geph_RNAi_; ** p=0.0063, WT vs SMA; *** p<0.0001, SMA vs SMA+Geph_RNAi_; OneWay ANOVA, Tukey’s *post hoc* test. “ns”: not significant. Relative expression of *nkcc1* and *kcc2* in medial L5 motor neurons **(F)** and lateral L5 motor neurons **(G)** in wild type (N=3) and SMA (N=3) mice. **(H)** Average life span in SMA mice injected with AAV9-GFP (as controls, N=9 mice) or with AAV9-Gephyrin_RNAi_-GFP (N=15 mice). **(I)** Body weight gain in wild type (blue, N=17 mice), wild type mice treated with Org25543 (cyan, N=7 mice), SMA mice (red, N=7 mice) and SMA mice treated with Org25543 (purple, N=9 mice).

